# A sphingomyelin-cholesterol complex regulates TLR4 activation in microglia

**DOI:** 10.64898/2026.09.10.750647

**Authors:** Cristina Antón-Barros, Beatriz Villarejo-Zori, Sara Carmona-Lorenzo, Juan Manuel Barbero-Paredes, Daniel González-Díaz, Mónica Uceda, Anna Fernàndez-Bernal, Manuel Portero-Otín, Iván Ayuso-Fernández, Cristina Galiana-Roselló, Francisco García-García, Elisa Navarro, Jorge Montesinos

**Affiliations:** Department of Biomedicine, Centro de Investigaciones Biológicas Margarita Salas. CSIC, Madrid, Spain; Escuela de Doctorado, Universidad Autónoma de Madrid, Madrid, Spain; Laser Confocal and Multidimensional in vivo Microscopy Unit, Centro de Investigaciones Biológicas Margarita Salas. CSIC, Madrid, Spain; Metabolic Pathophysiology Research Group, Department of Experimental Medicine, University of Lleida (UdL), Lleida, Spain; Biomedical Research Institute of Lleida - Dr. Pifarré Foundation, IRBLleida. Metabolic Pathophysiology Laboratory, Universitat de Lleida-Irblleida, Lleida, Spain; Department of Biotechnology, Centro de Investigaciones Biológicas Margarita Salas. CSIC, Madrid, Spain; Computational Biomedicine Laboratory, Principe Felipe Research Center (CIPF), Valencia, Spain; Department of Biochemistry and Molecular Biology, School of Medicine, Complutense University of Madrid, Madrid, Spain; CIBERNED, Network Center for Biomedical Research in Neurodegenerative Diseases, Madrid, Spain

**Author notes:** Correspondence to: Dr. Montesinos, Centro de Investigaciones Biológicas Margarita Salas-CSIC, C/Ramiro de Maeztu, 9. Madrid (Spain).

**Keywords:** Toll-like receptor 4, microglia, inflammation, SR-B1, cholesterol, sphingomyelin, lipidomics

## Abstract

Microglial activation in response to lipopolysaccharide (LPS) requires Toll-like receptor 4 (TLR4) redistribution into cholesterol- and sphingolipid-rich membrane domains, yet the lipid determinants of this process remain unclear. Here, we identify cholesterol accessibility as a key driver of TLR4-dependent microglial activation. We show that LPS increases cholesterol uptake and mobilization, whereas inhibition of intracellular cholesterol trafficking or the plasma membrane-to-ER cholesterol transporter Aster attenuates inflammatory responses. Scavenger receptor class B type 1 (SR-B1) mediates LPS-induced cholesterol uptake, and its inhibition suppresses TLR4 signaling, receptor recruitment to detergent-resistant membrane domains, and TLR4 endocytosis without altering bulk cellular cholesterol. Instead, SR-B1 inhibition remodels the sphingolipidome and expands the sphingomyelin-associated cholesterol pool, as detected by OlyA, while LPS promotes remodeling of this pool and increases accessible cholesterol detected by D4H. Manipulation of sphingomyelin-cholesterol interactions alters membrane properties and TLR4 trafficking, while stabilization of these complexes suppresses inflammatory activation. In vivo, LPS induces remodeling of sphingomyelin-associated cholesterol in microglia, whereas Alzheimer’s disease-related ApoE4/Trem2^R47H^ microglia show impaired cholesterol-pool remodeling and enhanced inflammatory responses. Together, these findings establish sphingolipid-dependent cholesterol accessibility as a key regulatory layer controlling TLR4 organization and microglial inflammatory activation, positioning SR-B1 as an upstream regulator of this lipid-signaling axis.

## Introduction

Microglia are the resident immune cells of the central nervous system and play essential roles in host defense, tissue homeostasis, and neuroinflammation^1^. In response to pathogens or danger signals, microglia activate pattern-recognition receptors such as Toll-like receptor 4 (TLR4), which detects lipopolysaccharide (LPS) and triggers signaling cascades that lead to cytokine production, nitric oxide synthesis, and phagocytic activation^2,3^. Although the downstream signaling pathways of TLR4 have been extensively characterized, increasing evidence indicates that receptor activation is not determined solely by ligand binding but also by the lipid organization of the plasma membrane in which signaling occurs^4–6^.

TLR4 signaling requires receptor redistribution into ordered membrane domains enriched in cholesterol and sphingolipids^7^, often referred to as lipid rafts^8,9^. These domains facilitate receptor clustering, adaptor recruitment, and downstream signaling^10^, while receptor internalization from the plasma membrane controls signal propagation and termination^11,12^. Consequently, the biophysical properties of membranes, such as fluidity, which are determined by lipid composition^13^, have emerged as important regulators of innate immune signaling. However, how membrane lipid remodeling is coordinated during microglial activation remains incompletely understood.

Cholesterol is a central determinant of membrane organization and immune receptor function. In addition to serving as a structural component of bilayers, cholesterol participates in domain formation, vesicular trafficking, and receptor nanoclustering^14–16^. Cellular cholesterol exists in distinct functional pools: an “accessible” fraction that can be trafficked to intracellular compartments and sensed metabolically, and a sphingomyelin-complexed fraction that is sequestered within the plasma membrane^17^. In parallel, perturbations in sphingolipid metabolism have been linked to immune activation, phagocytosis, and inflammatory signaling^18–20^, suggesting that coordinated regulation of sphingolipids and cholesterol may be required for efficient receptor function. The balance between these pools is increasingly recognized as a key regulator of cellular signaling processes, yet its relevance to innate immune receptor activation in microglia has not been fully explored.

Here, we investigated how cholesterol trafficking and membrane lipid organization control TLR4-dependent microglial activation. Using pharmacological perturbation, lipidomic profiling, membrane biophysical analyses, and in vivo validation, we show that the cholesterol transporter SR-B1 regulates inflammatory signaling by modulating the balance between accessible and sphingomyelin-bound cholesterol. Our findings identify sphingomyelin-cholesterol coupling as a key determinant of microglial activation and reveal membrane lipid organization as an upstream regulatory layer of innate immune signaling in the brain.

## Results

### Cholesterol trafficking supports LPS-induced microglial activation

Because cholesterol has been implicated in TLR4 signaling^21^, we first investigated whether extracellular cholesterol availability contributes to microglial inflammatory activation. BV2 microglial cells were cultured in lipoprotein-free (LPF) serum, which lacks exogenous cholesterol, and stimulated with LPS. Under LPF conditions, LPS-induced nitrite production was significantly reduced compared to cells maintained in 10% FBS (**Fig. 1A**), indicating that extracellular cholesterol availability supports the inflammatory response.

**Figure 1.**
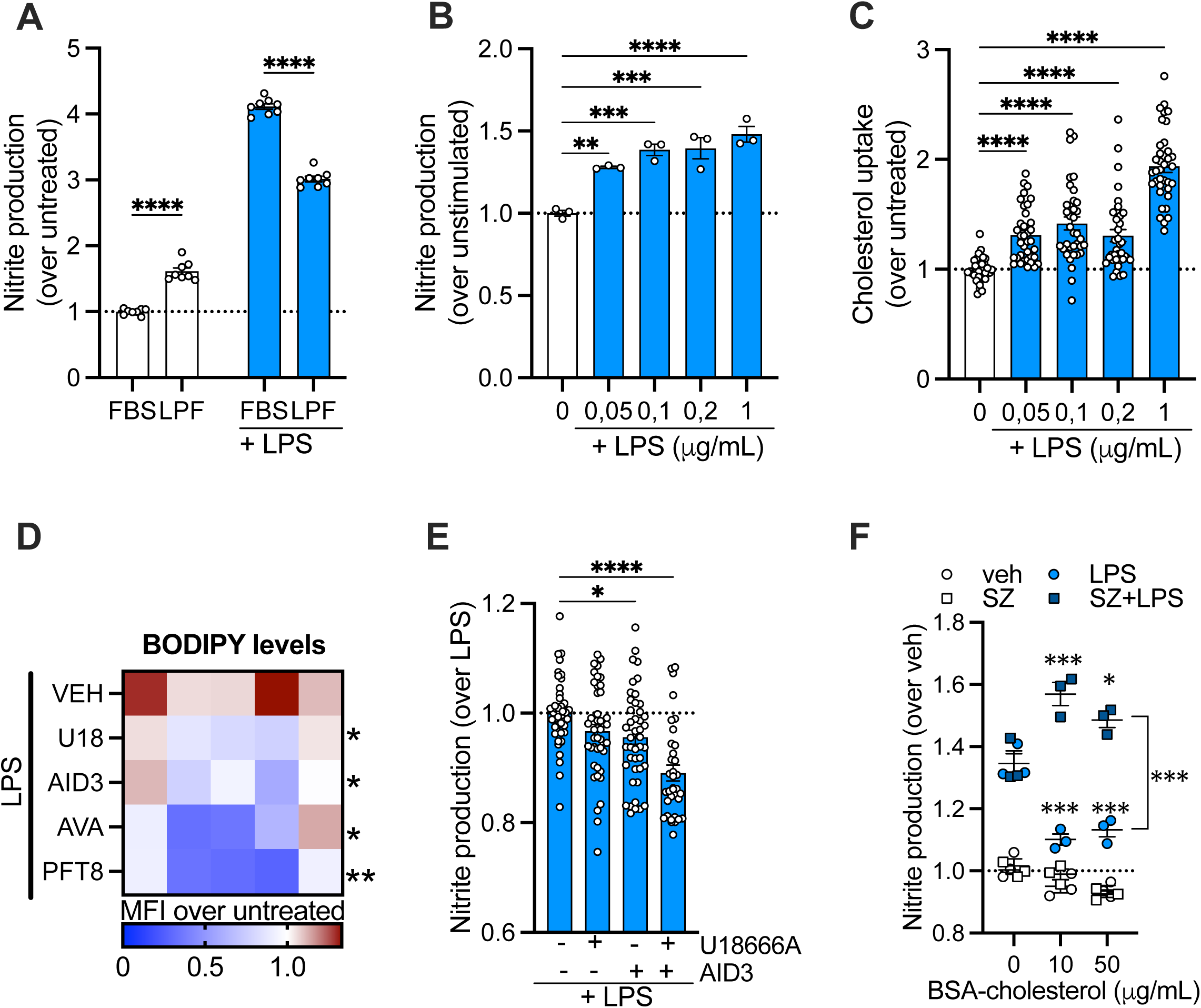
Cholesterol trafficking modulates LPS-induced inflammation. **A.** Nitrite production was measured in BV2 cells stimulated with LPS for 18h in the presence of 10% FBS or lipoprotein-free serum (LPF). **B-C.** BV2 cells were stimulated for 18h with the indicated concentrations of LPS, and nitrite production (**B**) and cholesterol uptake (**C**) were quantified. **D.** Lipid droplet accumulation was assessed by BODIPY staining in BV2 cells stimulated with LPS for 18h in the presence of U18666A (U18), AID3, avasimibe (AVA) and a combination of PF06424439 and T863 (PFT8). DMSO was used as vehicle (VEH). Data are shown as mean fluorescence intensity (MFI) relative to untreated control. **E.** Nitrite production was measured 18h after LPS stimulation in the presence of U18666A and/or AID3. **F.** BV2 cells were incubated for 24h with increasing concentrations of BSA-cholesterol in the presence or absence of the SOAT1 inhibitor Sandoz (SZ), followed by stimulation with LPS for 18h. Nitrite production was then assessed.

To further assess the role of cholesterol metabolism during inflammatory stimulation, we quantified cholesterol uptake using the fluorescent cholesterol analog NBD-cholesterol. As expected, LPS treatment increased nitrite production (**Fig. 1B**) and simultaneously enhanced cholesterol uptake in a dose-dependent manner (**Fig. 1C**). Similarly, IFNγ stimulation also promoted both cholesterol uptake and nitrite production, whereas treatment with IL1β or the TLR3 agonist poly(I:C) failed to induce comparable effects (**Fig. S1A-B**). These findings suggest that specific pro-inflammatory stimuli coordinately enhance cholesterol uptake.

To determine whether intracellular cholesterol trafficking contributes to the inflammatory response, cells were treated with U18666A, an inhibitor of vesicular and non-vesicular cholesterol transport, or AID-3, an inhibitor of the plasma membrane-to-ER cholesterol transporter Aster^22^. LPS stimulation induced the accumulation of neutral lipids, detected by BODIPY staining, consistent with increased cholesterol mobilization, esterification and lipid droplet formation (**Fig. 1D**). This effect was markedly reduced in the presence of either U18666A or AID-3, similarly to the inhibition observed with the SOAT1 inhibitor avasimibe, indicating impaired cholesterol delivery to the endoplasmic reticulum for esterification. We found similar results when using primary microglia (**Fig. S1C**). Importantly, inhibition of cholesterol trafficking attenuated the inflammatory response. AID-3 significantly reduced LPS-induced nitrite production, while U18666A produced a more modest effect (**Fig. 1E**). Combined inhibition further suppressed nitrite production, supporting a functional role for intracellular cholesterol mobilization in TLR4-mediated inflammatory signaling.

Finally, we investigated the impact of cholesterol overload on inflammatory response. Cells were incubated for 24 h with either BSA-complexed cholesterol (**Fig. 1F**) or MβCD-complexed cholesterol (**Fig. S1D**) prior to LPS stimulation. Cholesterol loading reduced LPS-induced nitrite production under both conditions. However, when cholesterol esterification was simultaneously blocked using the SOAT1 inhibitor Sandoz 58-035, nitrite production was significantly enhanced. These findings further support the idea that intracellular cholesterol mobilization, trafficking and esterification critically regulate inflammatory signaling in microglia.

### SR-B1 mediates LPS-induced cholesterol uptake in microglia

To identify the pathways responsible for cholesterol uptake in microglia, we first analyzed the expression of cholesterol transporters in publicly available RNA-seq datasets. Neonatal microglia, adult microglia, and the BV2 microglial cell line all expressed the cholesterol transporters *Ldlr*, *Cd36*, and *Scarb1*, with *Scarb1* showing the highest expression levels across developmental stages (**Fig. 2A**). We further validated the expression of these transporters in BV2 cells and primary microglia by qPCR, confirming robust expression of *Scarb1* and *Cd36*, with *Scarb1* predominating in BV2 cells and *Cd36* showing higher expression in primary microglia (**Fig. 2B**).

**Figure 2.**
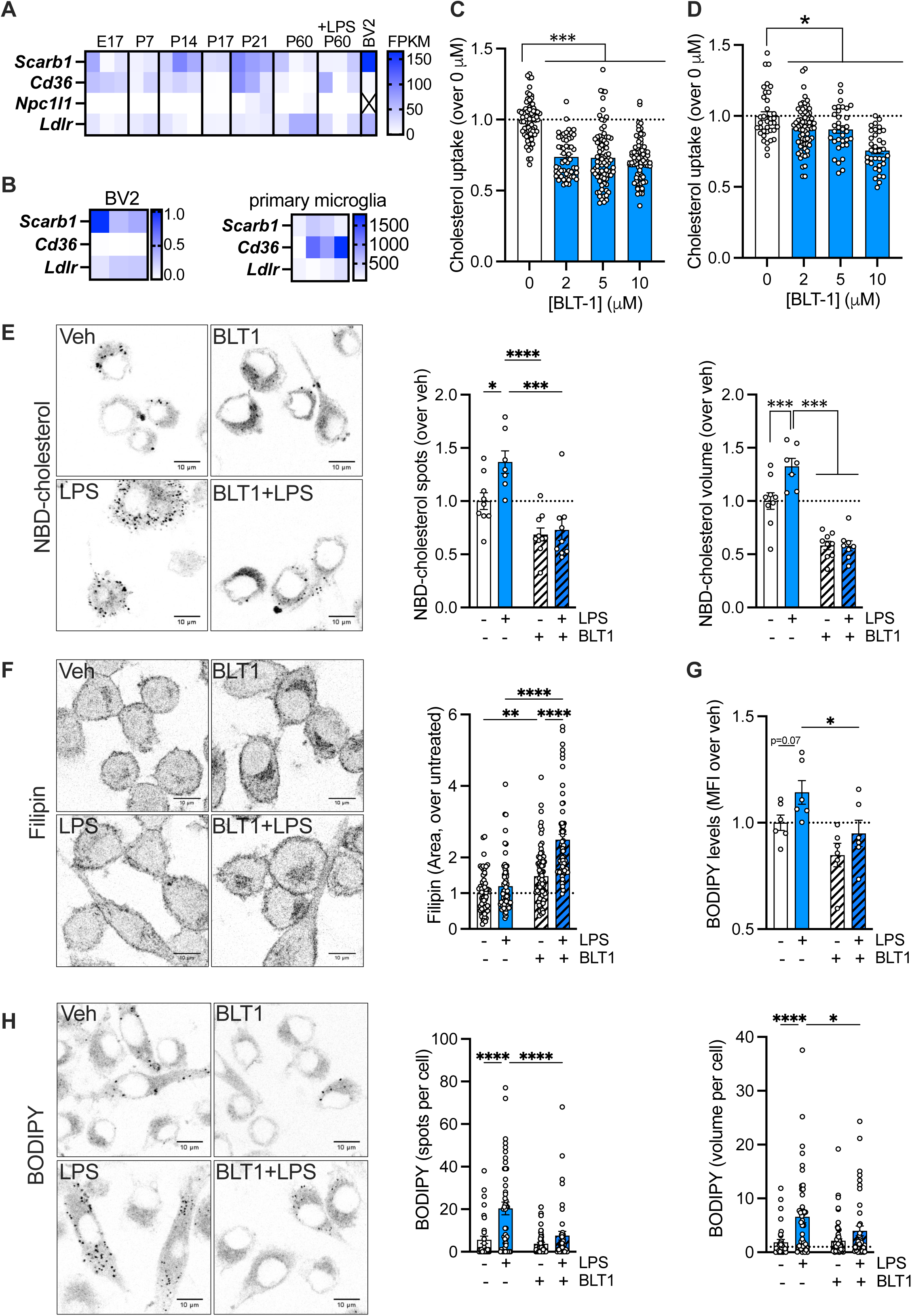
SR-B1 inhibition with BLT-1 alters cholesterol trafficking in microglia. **A.** Publicly available RNA-seq databases^66,67^ (<u>brainrnaseq.org</u>) were consulted to evaluate the expression levels of the indicated genes in microglia throughout mouse development. For BV2 gene expression, data was extracted from previous work^68^. **B.** Expression of *Scarb1*, *Cd36* and *Ldlr* was analyzed by RT-qPCR in BV2 cells and primary microglia. Data are shown as relative expression normalized to *18S*. **C-D.** Cholesterol uptake was determined by multipoint analysis of fluorescent NBD-cholesterol incorporation in the presence of increasing concentrations of the SR-B1 inhibitor BLT-1 in BV2 cells (**C**) and primary microglia (**D**) after 3h. **E.** BV2 cells were treated with 2 µM BLT1 for 18h and subsequently incubated with NBD-cholesterol for 3h in the presence or absence of LPS. NBD-cholesterol uptake was analyzed by confocal microscopy, and the number and volume of NBD-cholesterol spots were quantified. **F.** Filipin staining was used to detect cholesterol levels and distribution in BV2 cells pre-treated with 2 µM BLT1 for 18h and stimulated for 24h with LPS. **G-H.** Neutral lipids were detected by BODIPY staining and analyzed by flow cytometry (**G**) or visualized by confocal microscopy (**H**) in BV2 cells pre-treated with 2 µM BLT1 for 18h and stimulated for 24h with LPS. Image analysis was used to quantify BODIPY spots per cell and volume per cell.

We next examined the functional contribution of these transporters to cholesterol uptake. BV2 cells and primary microglia were treated with BLT-1 or SSO, inhibitors of SR-B1 and CD36^23,24^, respectively, and cholesterol uptake was quantified using NBD-cholesterol. In BV2 cells, inhibition of either SR-B1 or CD36 reduced cholesterol uptake, although BLT-1 produced the strongest effect at the lowest concentration tested (**Fig. 2C & S2A**). In contrast, CD36 inhibition with SSO failed to reduce cholesterol uptake in primary microglia (**Fig. S2B**), whereas BLT-1 treatment consistently suppressed uptake similarly to that observed in BV2 cells (**Fig. 2D**). Importantly, none of these treatments affected cell viability, as determined by MTT or crystal violet assay, except a slight reduction at 10 µM BLT1 (**Fig. S2C-F**). These findings identify SR-B1 as a major pathway mediating cholesterol uptake in microglia.

To determine whether SR-B1 contributes to cholesterol accumulation during inflammatory activation, BV2 cells were stimulated with LPS upon pretreatment with 2 µM BLT-1. Cholesterol uptake was assessed by quantifying NBD-cholesterol puncta number and volume (**Fig. 2E**), together with filipin staining for free cholesterol (**Fig. 2F**) and BODIPY staining for neutral lipids (**Fig. 2G-H**). BLT-1 markedly reduced the LPS-induced uptake of NBD-cholesterol and accumulation of neutral lipids. Analysis of filipin-stained cells revealed no significant changes following LPS stimulation alone, likely reflecting the rapid mobilization of cholesterol into cholesteryl esters. In contrast, BLT-1 treatment led to a significant accumulation of free cholesterol, particularly at the plasma membrane and within intracellular compartments, consistent with impaired cholesterol trafficking and esterification. Collectively, these findings indicate that SR-B1-mediated lipid uptake contributes substantially to cholesterol trafficking during microglial activation.

To further investigate the mechanism underlying these effects, we examined whether SR-B1 inhibition altered cholesterol efflux or endogenous cholesterol synthesis. BLT-1 treatment did not modify cholesterol efflux capacity (**Fig. S2G**) or the expression levels of *Abca1* and *Ch25h* (**Fig. S2H**), and no significant changes were detected in the expression of cholesterol biosynthetic genes *Hmgcr* and *Sqle* or the cholesterol transporters *Ldlr*, *Cd36* or *Scarb1* (**Fig. S2H**). Together, these findings indicate that SR-B1 primarily regulates cholesterol accumulation in LPS-treated microglia through modulation of cholesterol trafficking rather than altered synthesis or efflux pathways.

### SR-B1 inhibition suppresses inflammatory signaling and alters TLR4 membrane dynamics

To determine whether SR-B1-mediated cholesterol uptake contributes to microglial inflammatory activation, BV2 microglial cells were pretreated overnight with the SR-B1 inhibitor BLT-1 prior to LPS stimulation. Inhibition of SR-B1 markedly reduced LPS-induced nitrite release and IL-1β production (**Fig. 3A-B**), indicating that SR-B1 activity supports pro-inflammatory signaling. In addition, BLT-1 attenuated the increase in phagocytic activity induced by LPS stimulation (**Fig. 3C**), suggesting a broader impact on microglial activation programs.

**Figure 3.**
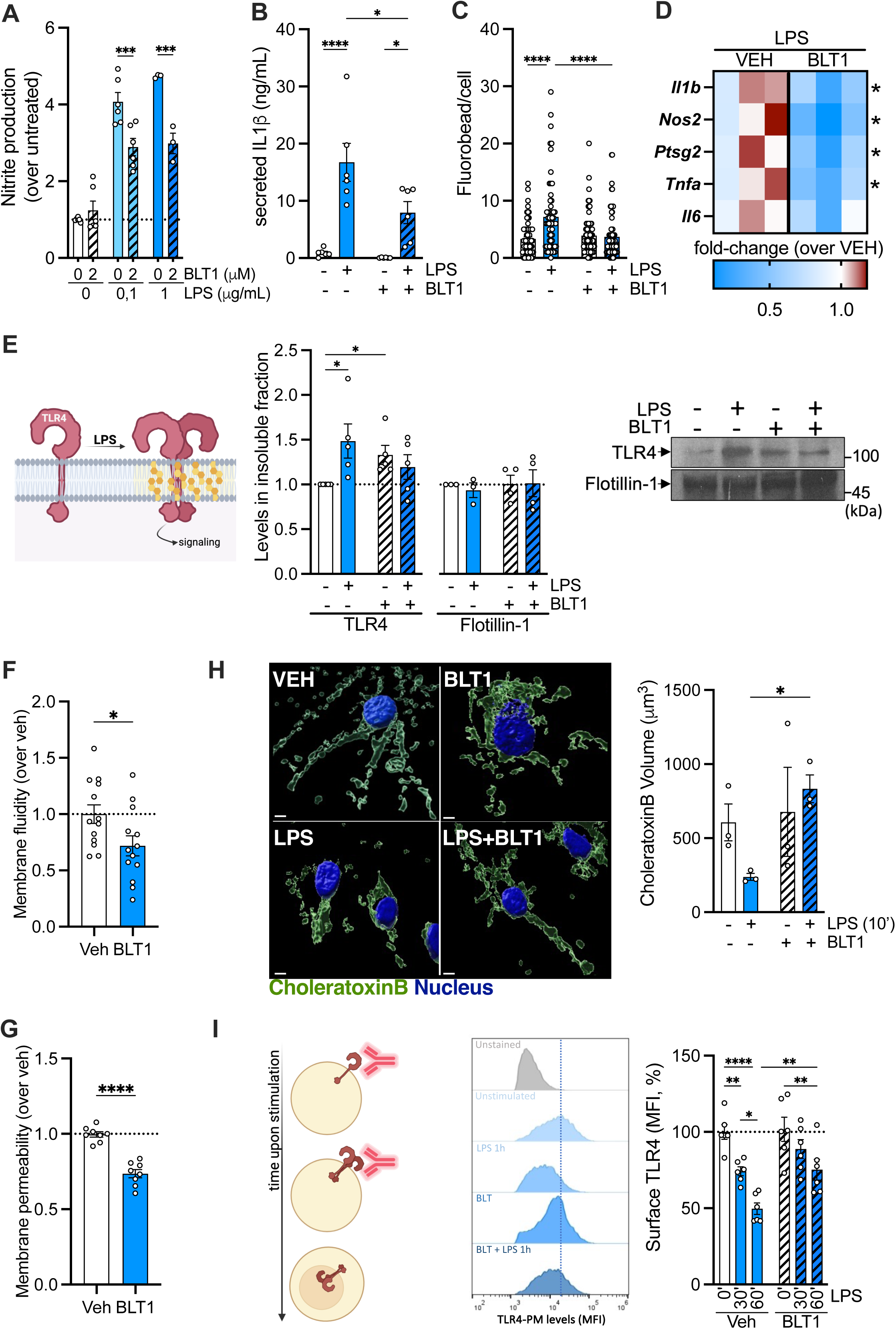
TLR4 inflammatory signaling is disrupted upon BLT-1 treatment. **A.** Nitrite production was measured in BV2 cells pre-treated for 18h with BLT1 and stimulated with the indicated concentrations of LPS for 24h. **B.** BV2 cells were treated for 18h with BLT-1 and then exposed to 1 µg/mL of LPS. 24h later, IL1β levels in the media were analyzed. **C.** Phagocytic activity was evaluated by confocal imaging of fluorescent microspheres (fluorobeads) uptake. **D.** Expression of the indicated pro-inflammatory genes was analyzed in primary microglia pre-treated with BLT-1 for 18h and then stimulated with LPS for 24h. **E.** TLR4 levels in detergent-insoluble fractions from BV2 cells pre-treated with BLT-1 and stimulated with LPS for 5min were analyzed by western blot. Flotillin-1 levels are shown as a control. **F-G.** Membrane fluidity (**F**) and membrane permeability (**G**) were assessed in BV2 cells following 18h of BLT-1 treatment. **H.** Lipid raft-associated GM1 was detected with fluorescently labeled cholera toxin B (green) in primary microglia pre-treated with BLT-1 for 18h and stimulated with LPS for 10min. Nuclei were stained with DAPI. Confocal imaging and IMARIS analysis were used to quantify total cholera toxin B volume. **I.** Surface TLR4 levels were analyzed by immunostaining and flow cytometry in BV2 cells pre-treated with BLT-1 and stimulated with LPS for the indicated times. Mean fluorescence intensity is expressed as percentage relative to unstimulated control.

We next validated these findings in primary microglia. Consistent with the effects observed in BV2 cells, BLT-1 significantly reduced LPS-induced expression of the inflammatory genes *Il1b*, *Nos2*, *Ptgs2*, and *Tnfa* (**Fig. 3D**), as well as IL-1β secretion, ROS generation, and phagocytic activity (**Fig. S3A-C**). In contrast, inhibition of CD36 with SSO failed to produce comparable effects (**Fig. S3D-E**), consistent with its limited contribution to cholesterol uptake in primary microglia.

TLR4 activation is known to involve receptor redistribution into cholesterol-rich detergent-resistant membrane domains followed by receptor endocytosis^7,25^. To determine whether SR-B1 inhibition affected these processes, Triton X-100–insoluble membrane fractions were isolated following LPS stimulation and TLR4 levels were quantified. BLT-1 pretreatment reduced the amount of TLR4 recovered in detergent-resistant membrane fractions after LPS exposure (**Fig. 3E**), indicating impaired recruitment of TLR4 into signaling-competent membrane domains.

Indeed, SR-B1 inhibition altered membrane biophysical properties associated with receptor signaling, such as reduced plasma membrane fluidity (**Fig. 3F**) and membrane permeability (**Fig. 3G**), consistent with altered membrane lipid organization. Appropriate fluidity controls are included in **Fig. S3F**. To further examine lipid raft organization, the raft-associated lipid GM1 was detected using cholera toxin B in primary microglia. Total GM1 levels at the plasma membrane showed a small increase following BLT-1 treatment in BV2 cells (**Fig. S3G**), and, using primary microglia, high-resolution image analysis revealed profound alterations in GM1 spatial organization and clustering (**Fig. 3H**).

Because TLR4 internalization occurs following its recruitment to lipid rafts, we next assessed receptor endocytosis after LPS stimulation. BLT-1 treatment delayed LPS-induced TLR4 internalization in BV2 cells (**Fig. 3I**) and almost completely abolished receptor endocytosis in primary microglia (**Fig. S3H**). Notably, BLT-1-treated primary microglia also displayed reduced basal surface TLR4 levels compared to untreated cells. Together, these findings indicate that SR-B1 inhibition alters membrane biophysical organization and impairs TLR4 redistribution into signaling-competent membrane domains, thereby suppressing downstream inflammatory signaling in microglia.

### SR-B1 inhibition remodels the sphingolipidome without altering bulk cholesterol

Because our data indicated alterations in membrane organization and cholesterol-rich domains, we next quantified cellular cholesterol levels following LPS and BLT-1 treatment (**Fig. S4A-C**). Surprisingly, neither total cholesterol nor cholesteryl ester (CE) content differed among the tested conditions. Likewise, cholesterol levels within detergent-resistant fractions remained unchanged (**Fig. S4D**). Thus, the changes in membrane organization associated with LPS stimulation and SR-B1 inhibition do not appear to reflect changes in bulk cellular cholesterol abundance.

We therefore examined the broader lipidomic response to LPS and BLT-1 in BV2 cells (**Table S1&S2**). LPS induced bidirectional remodeling of glycerophospholipid and sphingolipid species, together with TAG accumulation, whereas BLT-1 produced a more limited response, primarily characterized by increases in specific sphingolipid species and acylcarnitines (**Fig. 4A & S4E**). The combined LPS+BLT condition showed the largest number of differentially abundant species compared with control. Although a substantial component of the LPS-induced signature was preserved following BLT-1 pretreatment, with 64 species significantly altered in both conditions and strong concordance in effect size (r = 0.81; **Fig. S4F-G**), BLT-1 additionally altered the abundance of specific lipid species (**Fig. S4H-I**). These changes were particularly evident among sphingolipids as well as acylcarnitines, suggesting that SR-B1 inhibition substantially modifies the lipidomic response to inflammatory stimulation.

**Figure 4.**
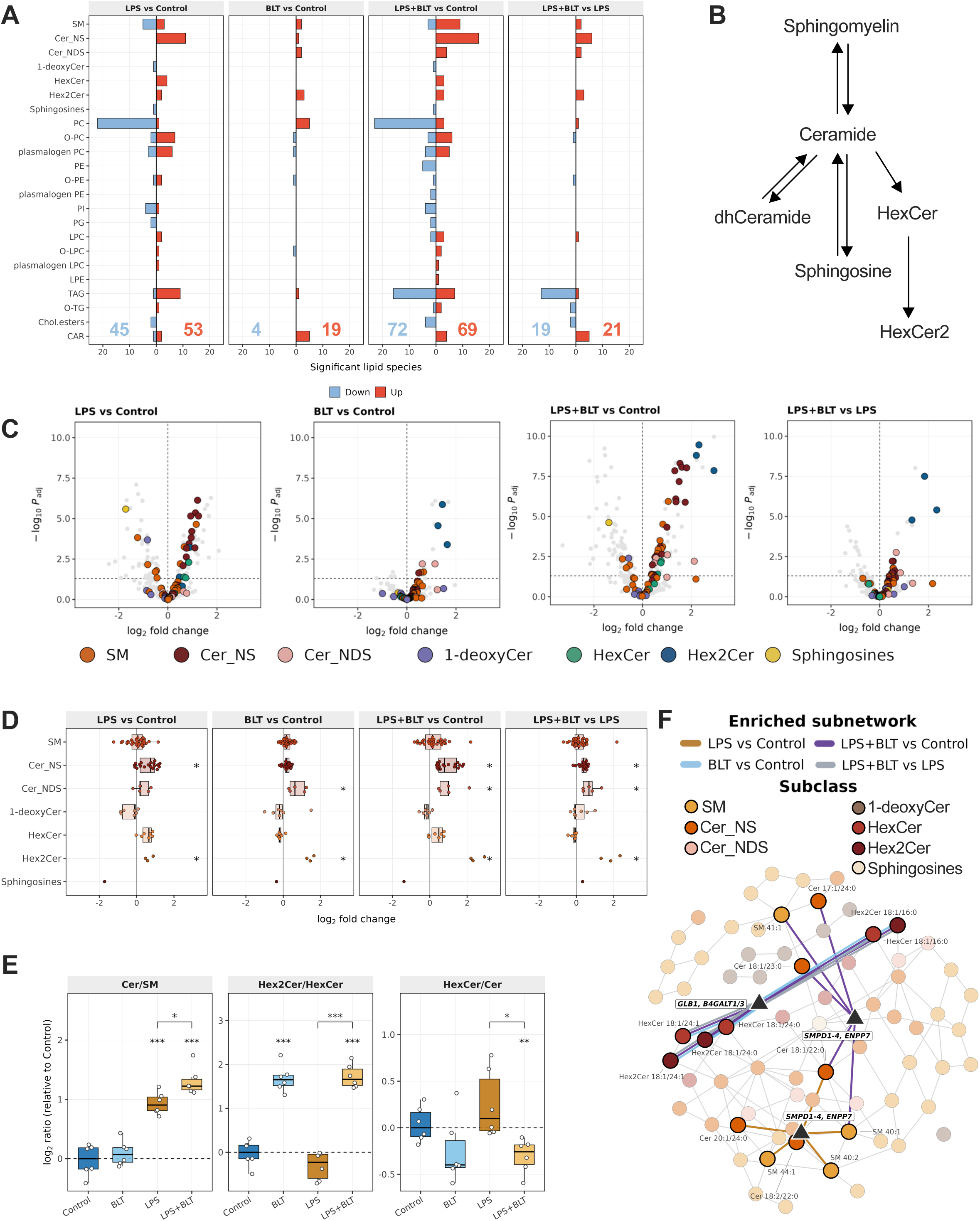
BLT-1 rewires the sphingolipidome. **A.** Lipid extracts of BV2 cells pre-treated with BLT-1 for 18h and stimulated with LPS for 24h were used for lipidomic analysis and differentially abundant lipid species were calculated (n=6). Bars indicate the number of lipid species significantly increased (up) or decreased (down) per subclass and contrast. Only species passing FDR < 0.05 are counted. Coloured numbers indicate the total count of significant lipid species in each direction. **B.** Schematic representation of sphingolipid biosynthesis highlighting the metabolic relationships between sphingomyelin (SM), ceramide (Cer; Cer_DS), dihydroceramide (dhCer; Cer_NDS), sphingosine and hexosylceramide species (HexCer and HexCer2). **C.** Volcano plots of the sphingolipid panel. All sphingolipid species are coloured by subclass irrespective of significance; the remaining species are shown in grey. Dashed lines mark zero fold change and FDR = 0.05. **D.** Species-level log2 fold changes of the sphingolipids. Each point is one lipid species and the box shows the dispersion within the subclass; boxes are drawn only for subclasses with five or more species. An asterisk marks subclasses reaching FDR < 0.05 in the class-level test, which is a self-contained rotation test (fry) treating the subclass as a set and accounting for the correlation among its species; the position of the box reflects the per-sample summary centred by species, not the individual contrasts. Ten species with missing values are excluded from the class-level test but retained in the species-level analysis. Subclasses with a single species are shown for completeness, but they have no class-level test. **E.** Sphingolipid class ratios.The panels are centred on the median of the control group and only the shift between groups is interpretable. Asterisks indicate FDR < 0.05, 0.01 and 0.001 relative to the control group. Each panel has its own y scale. **F.** LINEX-enriched subnetwork showing reaction relationships among sphingolipid species across the four experimental contrasts. Nodes are colored by lipid subclass, and colored edges indicate contrast-specific enriched reactions, mainly involving SM-Cer and HexCer-Hex2Cer transitions. Edges represent composition-compatible pathway relationships and should not be interpreted as definitive one-to-one molecular conversions for incompletely resolved species..

Because sphingolipids represented a prominent component of the BLT-1-associated response, we next examined this lipid class in greater detail. Our lipidomic analysis resolved several sphingolipid classes, including sphingomyelins (SM), canonical ceramides (Cer_NS), dihydroceramides (Cer_NDS), monohexosylceramides (HexCer), dihexosylceramides (Hex2Cer), 1-deoxyceramides (1-deoxyCer), and sphingosine (**Fig. 4B**). LPS predominantly increased Cer_NS species, whereas BLT-1 was more strongly associated with Cer_NDS and Hex2Cer. Notably, LPS stimulation following BLT-1 pretreatment resulted in marked remodeling of SM, Cer and Hex2Cer species, encompassing both Cer_NS and Cer_NDS (**Fig. 4C-D**). Analysis of class ratios (**Fig. 4E**) further revealed distinct effects of LPS and BLT-1 on relationships within the sphingolipid network. LPS increased the summed Cer/SM abundance ratio, whereas BLT-1 markedly increased the Hex2Cer/HexCer ratio both under basal conditions and following LPS stimulation. Together, these findings indicate that inflammatory stimulation and SR-B1 inhibition differentially remodel interconnected sphingolipid pools.

To further resolve these changes within the sphingolipid reaction network, we performed LINEX2-based substrate-product subnetwork enrichment (**Table S3**). LPS and BLT-1 showed distinct enriched reaction patterns (**Fig. 4F & S4J**). In LPS-treated cells, the predominant reaction subnetwork was centered on the SM↔Cer axis, and concentrated among long-chain SM and Cer species containing approximately 40-44 total carbons. In contrast, BLT-1 preferentially enriched relationships between HexCer and Hex2Cer, accompanied by increases in Cer_NDS and Hex2Cer species, particularly among species containing 42-43 total carbons. The combined LPS+BLT condition incorporated features of both responses, retaining the SM-Cer-associated signature observed with LPS while displaying the HexCer-Hex2Cer relationships characteristic of BLT-1. Consistently, direct comparison of LPS+BLT versus LPS identified HexCer↔Hex2Cer as the principal enriched subnetwork associated with BLT-1 under inflammatory conditions. These LINEX-derived patterns provide a molecular network-level interpretation of the lipidomic changes, suggesting that SR-B1 inhibition alters the remodeling of the sphingolipid network during inflammatory stimulation.

We next examined whether lipidomic remodelling involved structural features relevant to membrane organisation. LION enrichment revealed distinct treatment-dependent profiles (**Fig. S4K**). LPS was primarily characterised by enrichment of saturated species and species with a low degree of unsaturation, whereas BLT-1 showed enrichment of long- and very-long-chain sphingolipid species, particularly C24-containing ones, together with negative enrichment of polyunsaturated species, plasmalogens and several C18-C20-containing species. The two treatments therefore produced qualitatively different structural remodelling rather than a common shift along a single physicochemical axis. Terms in the biophysical branch - bilayer thickness, transition temperature, lateral diffusion and intrinsic curvature - were not interpreted as membrane-property changes because each was annotated predominantly within glycerophospholipids and was strongly influenced by lipid-class composition. Under BLT-1, for instance, the intrinsic-curvature terms were dominated by different glycerophospholipid subclasses, such that their apparent redistribution largely reflected the divergent behaviour of those classes. Nevertheless, the BLT-1-associated enrichment of C24 sphingolipids together with the reduced representation of polyunsaturated species and plasmalogens is compatible with tighter lipid packing and is consistent with the experimentally observed lower membrane fluidity relative to LPS.

Together, these findings demonstrate that SR-B1 inhibition does not alter bulk cellular cholesterol abundance but instead induces selective remodeling of the sphingolipidome, particularly within the Cer and glycosphingolipid branches. Moreover, BLT-1 modifies the lipidomic response to LPS by introducing a distinct sphingolipid signature, providing a potential lipidomic basis for the altered membrane organization and signaling associated with SR-B1 inhibition.

### Sphingomyelin-cholesterol complexes coordinate microglial response/ BLT-1 expands the sphingomyelin-cholesterol complex pool

SM forms stable complexes with cholesterol in cellular membranes, generating a biophysical pool of cholesterol that is considered “inaccessible” for trafficking and signaling. In contrast, “accessible” cholesterol is more dynamically available for transport to the endoplasmic reticulum, where it contributes to esterification and lipid remodeling^26^. Given that BLT-1 treatment remodelled SM and ceramide classes, we hypothesized that SR-B1 inhibition may alter the balance between these two cholesterol pools by expanding SM-cholesterol complexes without necessarily changing total cholesterol content.

To directly test this, we quantified SM-cholesterol complexes using the SM-binding probe OlyA^27^ and measured accessible cholesterol using the D4H-mCherry sensor^28^. We first validated the specificity of both approaches in BV2 microglia. OlyA detection was optimized using a His-tagged OlyA construct followed by fluorescent anti-His labeling (**Fig. 5A**). This system reliably reported SM-cholesterol complex abundance, as OlyA signal increased following cholesterol loading with MβCD-cholesterol and decreased after cholesterol extraction with MβCD or enzymatic disruption of SM-cholesterol complexes using sphingomyelinase (SMase) (**Fig. S5A**). In parallel, D4H-mCherry accurately reported changes in accessible cholesterol, increasing upon cholesterol loading and decreasing upon extraction (**Fig. S5B**).

**Figure 5.**
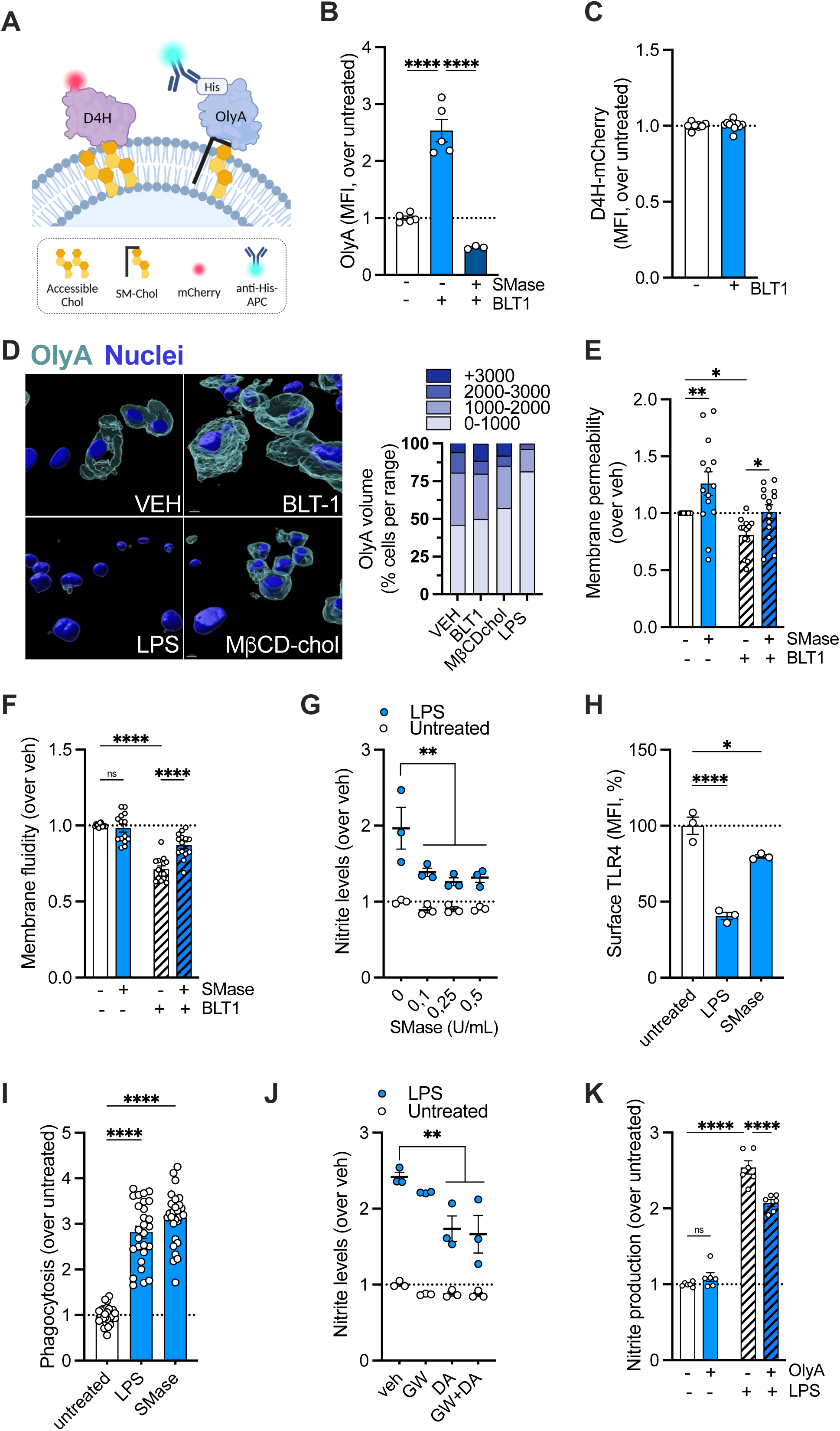
SM-cholesterol complexes regulate LPS-induced inflammation. **A.** SM-bound cholesterol was detected by fluorescent OlyA-His staining while accessible cholesterol was assessed using the fluorescent D4H-mCherry. **B.** OlyA staining increased after 18h of BLT-1 treatment while incubation with 0,5 U/mL sphingomyelinase (SMase) for 2h reduced OlyA signal. **C.** Accessible cholesterol levels, measured by D4H-mCherry fluorescence, were unaltered following 18h incubation with BLT-1. **D.** Primary microglia were treated with LPS or BLT-1 and analyzed for membrane SM:cholesterol complexes using OlyA staining. Cells incubated for 30min with 60 µg/mL MβCD-cholesterol were used as a positive control. OlyA signal was acquired by confocal microscopy and quantified using Imaris-based 3D image analysis to obtain OlyA^+^ volume per cell. Individual cells were then binned according to OlyA volume (0-1000, 1000-2000, 2000-3000, and >3000 arbitrary volume units). The proportion of cells within each volume category was calculated and expressed as a percentage of total cells per condition. Distribution shifts between conditions were further assessed using cumulative distribution analysis and the Kolmogorov-Smirnov test. **E-F.** Incubation with 0,5 U/mL SMase for 2h reversed the effects of BLT-1 on membrane permeability (**E**) and membrane fluidity (**F**). **G.** BV2 cells were incubated with the indicated concentrations of SMase for 30min prior to LPS stimulation, and nitrite production was measured. **H.** Surface TLR4 levels were analyzed in BV2 cells exposed to LPS or 0,5 U/mL SMase for 1h. **I.** Phagocytosis was assessed by multiwell analysis of fluorobeads uptake following 3h of exposure to LPS or 0,5 U/mL SMase. **J.** BV2 cells were treated for 18h with the neutral SMase inhibitor GW4869 (GW) or the acid SMase inhibitor desipramine (DA), followed by LPS stimulation and measurement of nitrite production. **K.** BV2 cells were incubated with 500 µg/mL OlyA for 18h and LPS-induced nitrite production was subsequently measured.

Having validated this system, we next examined the impact of BLT-1 on membrane cholesterol organization. SR-B1 inhibition significantly increased OlyA staining without altering D4H-mCherry signal (**Fig. 5B-C**), indicating a selective expansion of the SM-cholesterol complex pool without affecting accessible cholesterol levels. Importantly, the increase in OlyA signal induced by BLT-1 was reversed by SMase treatment, confirming that it reflects SM-dependent cholesterol sequestration.

We then investigated how pro-inflammatory stimulation affects these cholesterol pools. In primary microglia, high-resolution imaging accompanied by Imaris-based quantification confirmed that BLT-1 increased OlyA volume (**Fig. 5D**), consistent with our flow cytometry results. LPS stimulation induced a dramatic reduction in OlyA signal (**Fig. 5D**) while increasing labeling of D4H-mCherry (**Fig. S5C**) indicating the mobilization of SM-complexed cholesterol into accessible cholesterol. This LPS-induced remodeling of SM-rich domains was also confirmed by flow cytometry (**Fig. S5D**).

To assess the functional relevance of SM-cholesterol complexes, we perturbed this pool pharmacologically. Exogenous SMase treatment, which disrupts SM-cholesterol interactions, restored membrane permeability and fluidity in BLT-1–treated cells (**Fig. 5E-F**). However, SMase did not restore LPS-induced nitrite production in cells treated with BLT-1 (**Fig. S5E**). Similarly, acute cholesterol extraction with MβCD had no effect on BLT-1 reduction of nitrite production (**Fig. S5F**).

Interestingly, SMase treatment itself modulated inflammatory signaling in a time-dependent manner. Short exposure (30 min pretreatment) reduced LPS-induced nitrite production (**Fig. 5G**), while prolonged pretreatment (24 h) had no significant effect (**Fig. S5G**). This reduction was associated with increased TLR4 endocytosis and enhanced phagocytic activity (**Fig. 5H-I**), suggesting that acute disruption of SM-cholesterol organization facilitates receptor redistribution and endocytic processes. These results support the idea that inflammatory outcomes depend on the dynamic balance between accessible and SM-sequestered cholesterol pools rather than total cholesterol levels.

To further assess the contribution of endogenous SM turnover, we inhibited sphingomyelinase activity using the neutral SMase inhibitor GW4869 or the acid SMase inhibitor desipramine. Acute co-treatment with LPS had minimal effects on nitrite production (**Fig. S5H**), whereas prolonged pretreatment reduced LPS-induced inflammatory responses, particularly with desipramine (**Fig. 5J**). These effects were confirmed in primary microglia, where desipramine pretreatment reduced nitrite release, ROS levels, and IL1β production upon LPS stimulation (**Fig. S5I-K**).

We next exploited OlyA itself as a functional probe. High-dose overnight OlyA treatment, which stabilizes and sequesters SM-cholesterol complexes, significantly reduced LPS-induced nitrite production, phenocopying the effect of BLT-1 (**Fig. 5K**). Together, these results indicate that stabilization of the SM-cholesterol pool suppresses inflammatory activation, highlighting the functional importance of cholesterol accessibility in TLR4 signaling.

Finally, we examined whether this mechanism operates in vivo. Using Tmem119-GFP reporter mice subjected to systemic LPS challenge (**Fig. 6A**), we found that LPS significantly reduced OlyA staining in GFP⁺ microglia from male mice (**Fig. 6B**), with a similar but non-significant trend in females (**Fig. 6C**). In contrast, D4H-mCherry labeling remained unchanged (**Fig. 6D**), indicating selective remodeling of SM-cholesterol complexes during neuroinflammatory activation in vivo. In a disease-relevant context, we compared OlyA and D4H staining upon LPS stimulation of microglia from wild-type mice and mice carrying Alzheimer’s disease-associated ApoE4 and Trem2^R47H^ variants (**Fig. 6E-F**). Upon LPS stimulation, WT microglia exhibited a time-dependent reduction in OlyA staining while increasing D4H staining. In contrast, ApoE4/Trem2^R47H^ microglia showed no changes in the levels of either OlyA or D4H staining after stimulation, consistent with a chronic disruption of SM-cholesterol remodeling in response to LPS. Notably, these cells also displayed increased LPS-induced IL1β production (**Fig. 6G**), linking altered cholesterol accessibility to heightened inflammatory sensitivity.

**Figure 6.**
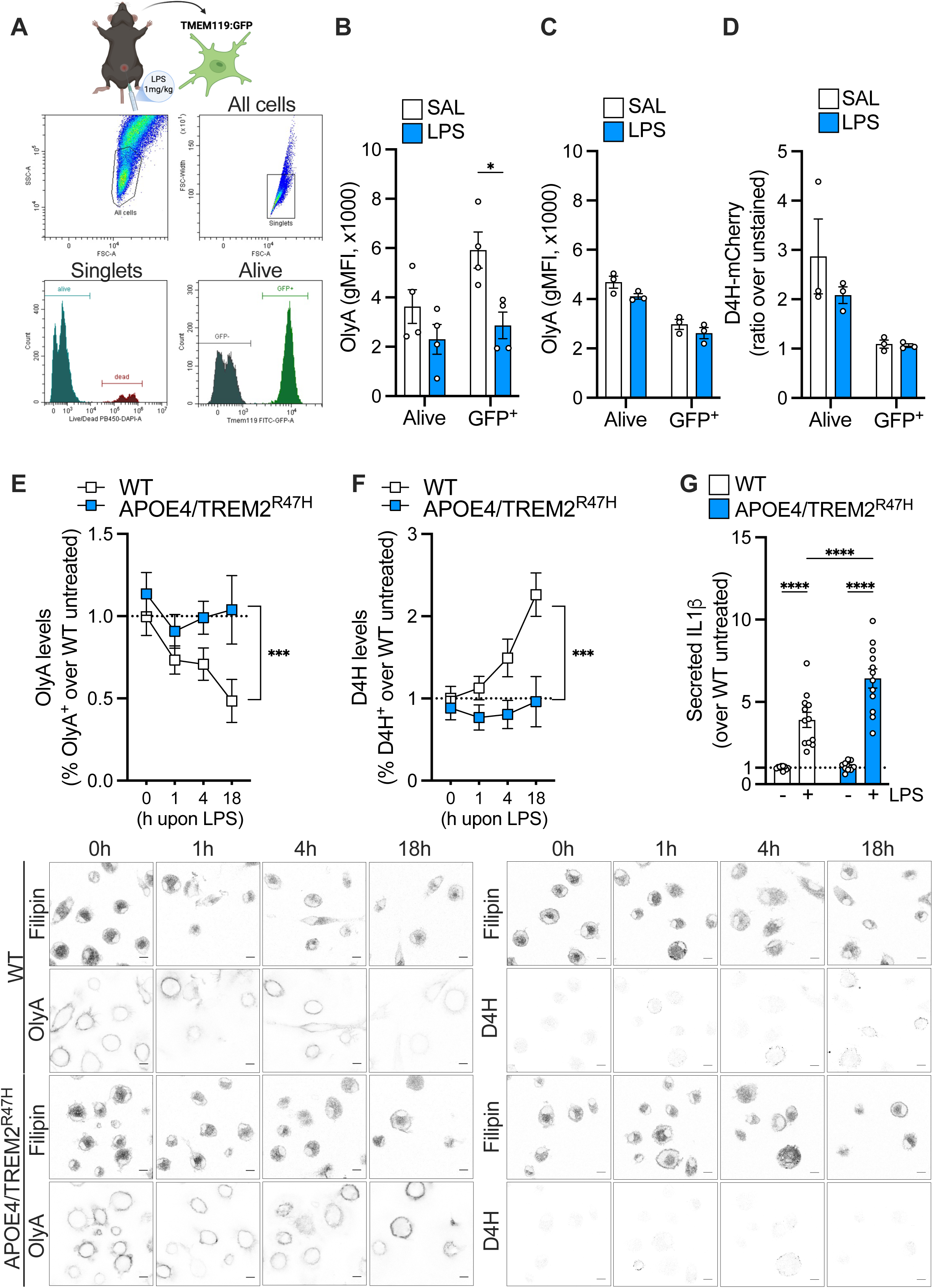
Altered SM-cholesterol complex levels are detected in a septic mouse model and in APOE4/TREM2^R47H^ microglia. **A.** Gating strategy used to quantify OlyA and D4H levels in viable microglia (DAPI^-^ Tmem119:GFP^+^) isolated from mouse brains following intraperitoneal injection of 1mg/kg LPS. **B-C.** Levels of SM-bound cholesterol, detected by OlyA staining, were analyzed in viable microglia (DAPI^-^ GFP+) and total viable cells (DAPI^-^) from saline (SAL)- and LPS-treated (**B**) male and (**C**) female mice. **D.** Accessible cholesterol levels were determined by D4H-mCherry staining in viable microglia (DAPI^-^ GFP+) and total viable cells (DAPI^-^) from saline (SAL)- and LPS-treated male mice. **E-F.** Primary microglia isolated from WT and APOE4/TREM2^R47H^ pups were cultured and stimulated with LPS for 0h, 1h, 4h or 18h and OlyA staining (**E**) and D4H staining (**F**) were then analyzed. For each condition, 39-105 individual cells were analyzed. Representative images for each condition are shown. **G**. The levels of IL1β were measured in the media upon 18h stimulation of LPS in WT and APOE4/TREM2^R47H^ microglia.

## Discussion

Collectively, our findings support a model in which SR-B1-mediated lipid flux regulates microglial inflammatory signaling by controlling membrane cholesterol accessibility. Using complementary cholesterol sensors, we show that LPS stimulation and SR-B1 inhibition redistribute cholesterol between accessible and inaccessible pools without substantially altering total cellular cholesterol. SR-B1 inhibition increases cholesterol association with sphingomyelin-rich membranes, shifting cholesterol toward an inaccessible pool. This redistribution is accompanied by reduced membrane fluidity, impaired TLR4 recruitment to detergent-resistant membrane domains, delayed receptor endocytosis, and attenuated inflammatory signaling. Importantly, manipulating sphingomyelin turnover with sphingomyelinase reverses these changes and restores inflammatory signaling, providing direct evidence that sphingomyelin-dependent cholesterol sequestration is a key determinant of TLR4 activity.

TLR4 signaling depends on receptor partitioning into cholesterol- and sphingolipid-enriched membrane domains, where clustering with CD14 and MD-2 facilitates MyD88-dependent NF-κB and MAPK activation^10,29,30^. Conversely, cholesterol depletion disrupts membrane-domain organization, impairs TLR4 clustering, and attenuates cytokine production^31^. Our data extend this model by showing that TLR4 signaling depends not simply on membrane cholesterol abundance, but on its partitioning between accessible and inaccessible pools.

This distinction is particularly relevant because sphingomyelin can sequester cholesterol into tightly packed complexes, limiting its accessibility and regulating its intracellular trafficking^32^. In our system, increased sphingomyelin following SR-B1 inhibition was associated with reduced cholesterol accessibility, membrane fluidity, and TLR4 recruitment and trafficking, whereas sphingomyelin depletion restored cholesterol accessibility and inflammatory signaling. Thus, sphingomyelin appears to actively determine the fraction of membrane cholesterol available for dynamic redistribution and signaling-related processes.

Cholesterol accessibility may also influence TLR4 through direct lipid-protein interactions. A cluster of cholesterol-binding CARC–CRAC–CARC motifs has been identified in the membrane-proximal intracellular region of TLR4, immediately downstream of the transmembrane domain and preceding the TIR domain^21^. This region overlaps a conserved hydrophobic sequence required for TLR4 activation and ligand-independent oligomerization^33^. Although direct cholesterol binding has not been demonstrated, these observations raise the possibility that local cholesterol accessibility could influence TLR4 conformation or oligomerization in addition to its membrane partitioning.

Our findings are further consistent with studies linking intracellular cholesterol trafficking to TLR4-dependent inflammation. Aster proteins transfer cholesterol from the plasma membrane to the endoplasmic reticulum thus regulating plasma membrane cholesterol levels; accordingly, Aster inhibition reduced LPS-induced TLR4 activation. Similarly, ABCA1-deficient macrophages accumulate free cholesterol and exhibit enhanced LPS responses^34^, together with increased cholesterol accumulation in lipid rafts and TLR recruitment to these domains^35^. Disruption of cholesterol-rich rafts attenuates this hyper-responsiveness^36^. These studies support a role for cholesterol delivery, efflux, and membrane organization in TLR signaling and, together with our findings, suggest that the balance between accessible and inaccessible cholesterol is a critical determinant of receptor activity.

Within this framework, SR-B1 emerges as an upstream regulator of membrane cholesterol organization. Although traditionally viewed as a lipoprotein receptor involved in cholesterol uptake and efflux during reverse cholesterol transport^37,38^, SR-B1 also regulates innate immune responses. SR-B1-deficient mice exhibit hyperinflammatory responses^39^, whereas SR-B1 overexpression attenuates inflammation^40^, without corresponding changes in total cellular cholesterol, suggesting that cholesterol distribution rather than abundance underlies these effects. Consistent with this, our data indicate that SR-B1 regulates inflammatory signaling primarily through membrane lipid organization and cholesterol accessibility.

This mechanism may involve SR-B1-dependent lipid flux across the plasma membrane. BLT-1 irreversibly blocks the hydrophobic channel of SR-B1^41^, potentially perturbing lipid flux and promoting compensatory sphingolipid remodeling. Indeed, SR-B1 inhibition preferentially altered long- and very-long-chain sphingomyelin species, an environment that may favor sphingomyelin-cholesterol complex formation and shift cholesterol toward an inaccessible pool. We therefore propose that SR-B1-dependent lipid flux maintains a dynamic equilibrium between accessible and sphingomyelin-sequestered cholesterol, whereas SR-B1 inhibition promotes cholesterol sequestration within sphingomyelin-rich domains and consequently alters membrane organization and TLR4 signaling.

Sphingolipid metabolism represents an additional regulatory layer linking cholesterol accessibility to inflammation. Cholesterol and sphingomyelin are reciprocally regulated: cholesterol availability influences very-long-chain sphingomyelin synthesis and trafficking, whereas sphingomyelin abundance determines cholesterol partitioning within membranes^42^. Genetic or pharmacological manipulation of sphingolipid metabolism modulates TLR4-dependent responses in macrophages, with effects depending on the pathway affected^20^. Consistent with this, our lipidomic data suggest that redirecting ceramide metabolism toward glucosylceramide rather than sphingomyelin synthesis is associated with impaired TLR4 signaling. Accordingly, macrophages deficient in serine palmitoyltransferase or sphingomyelin synthases display reduced TLR4/MD-2 surface recruitment and LPS-induced NF-κB activation, together with altered cholesterol efflux^43–45^.

The relevance of the sphingomyelin-cholesterol complex extends to neuroinflammation in vivo. Following systemic LPS challenge, reduced OlyA staining in microglia indicates remodeling or depletion of sphingomyelin-associated cholesterol, consistent with altered membrane cholesterol accessibility during inflammatory activation. Similarly, changes in OlyA and D4H staining in ApoE4/Trem2^R47H^ microglia after LPS challenge suggest that disease-associated microglial states involve altered sphingomyelin-dependent cholesterol organization. ApoE4 microglia exhibit increased lipid and lipid-droplet accumulation, altered morphology, and elevated cytokine and chemokine production^46,47^. The AD-associated TREM2^R47H^ loss-of-function variant is likewise associated with impaired lipid handling, increased cholesteryl ester accumulation, and defective clearance of myelin-derived cholesterol following lipid challenge^48,49^, whereas higher TREM2 expression is associated with enhanced cholesterol metabolism and increased phagocytic activity^50^. Together, these observations support a broader link between altered microglial cholesterol accessibility and disease-associated inflammatory states.

Overall, our findings position cholesterol accessibility as a central regulatory determinant of microglial immune signaling and identify SR-B1–dependent sphingolipid remodeling as an upstream mechanism controlling this process. TLR4 signaling appears to depend not on total cellular cholesterol, but on its partitioning between accessible and sphingomyelin-sequestered pools. By controlling this balance, SR-B1 regulates membrane organization, TLR4 trafficking, and inflammatory signaling. This framework explains how changes in lipid composition can profoundly alter innate immune responses without measurable changes in bulk cholesterol and suggests that targeting cholesterol accessibility or its sphingolipid-dependent regulation could provide a more selective strategy for modulating microglial inflammation.

## Material and Methods

### Animals and systemic acute LPS challenge

Wild-type (WT) C57BL/6 mice, Tmem119-GFP reporter mice and APOE4/Trem2^R47H^ mice on a C57BL/6 background were used. Mice were housed in groups of 4-5 in standard cages with free access to water and food under controlled environmental conditions at the CIB animal facility. To establish the systemic acute LPS challenge model, Tmem119-GFP mice received a single intraperitoneal injection of 1 mg/kg LPS (0,9% saline, 1mg/mL). Control mice were injected with equivalent volumes of physiological saline. Mice were sacrificed 18 h post-injection by cervical dislocation. Brains were immediately extracted and processed for flow cytometry experiments. All animal experiments were conducted following European Union guidelines and approved by the CSIC ethical review committee and the Comunidad de Madrid (PROEX224.0/24).

### Primary microglia isolation

Mixed glial cultures were prepared from postnatal day (P)0-P5 mouse brains (WT or APOE4/Trem2^R47H^). Pups were sacrificed by rapid decapitation, the brains were removed, and meninges were carefully peeled off. Cortices were mechanically dissociated by gentle trituration and the resulting suspension was filtered through a pre-wetted 70 µm cell strainer. Cells were centrifuged at 300 × g for 5 min and cultured in DMEM F12 supplemented with 10% FBS and 5 ng/mL GM-CSF. Cultures were maintained at 37°C in 5% CO₂, with weekly medium changes. After 14 days in vitro, microglia were purified by orbital shaking of confluent mixed glial cultures at 150 rpm in an orbital shaker for 3-4 h, followed by collection of the supernatant and centrifugation. The microglial pellet was resuspended, counted, and plated onto poly-L-lysine–coated cultureware at 30,000-40,000 cells/cm² in complete medium containing GM-CSF, and allowed to recover for at least 48h prior to stimulation. Microglia purity (>99%) was tested by Iba-1 immunostaining.

### Microglia cell line

BV2 microglial-like cells were cultured in DMEM supplemented with 10% FBS and 1% of penicillin/streptomycin and maintained at 37°C in 5% CO₂. Cells were routinely tested for mycoplasma.

### Stimulation and treatments

To induce a pro-inflammatory response, cells were challenged with 1 µg/mL lipopolysaccharide (LPS, from *E. Coli* E26:O6), unless otherwise stated. In some experiments, cells were stimulated with 1 ng/mL IL1β, 20 ng/mL IFNγ, and 10 µg/mL poly(I:C). To inhibit cholesterol uptake, cells were pre-treated with 2 µM BLT-1 (SR-B1 inhibitor), sulfosuccinimidyl oleate (SSO, CD36 inhibitor), 1 µM U18666A (NPC1/Aster inhibitor), or 2 µM AID3 (Aster inhibitor). Cholesterol esterification was inhibited using 5 µM avasimibe or 20 µM Sandoz 58-035 (SOAT1 inhibitors), whereas DGAT1/2 activity was inhibited using a combination of PF-06424439 and T863 (10 µM each). Neutral and acid sphingomyelinases were inhibited with 5 µM GW4869 and 10 µM desipramine, respectively. For cholesterol manipulation, MβCD-cholesterol and BSA-cholesterol were used for cholesterol supplementation while MβCD was used to deplete cholesterol. BSA-cholesterol was prepared in-house as previously described^51^. Equal volumes of DMSO were used as vehicle control. All experimental treatments were performed in 5% FBS DMEM for BV2 cells and 1% FBS DMEM-F12 for primary microglia at the indicated concentrations and incubation times.

### Nitrite measurement

Nitrite production was measured by Griess reaction. Briefly, cell media was incubated for 30 min at 37°C with prewarmed NEDA (300 µg/mL) and DAPSONE (1 mg/mL with 9% of HCl 5N). Absorbance at 540 nm was then measured using the Varioskan multimode microplate reader.

### Cell viability measurements

For crystal violet assay, cells were fixed with glutaraldehyde 1% and incubated with Crystal violet 0,1% for 30 min at r.t. After removing the dye excess, cells were dried overnight. Fixed crystal violet in the cells was resuspended with acetic acid 10% and absorbance was measured at 590 nm in the Varioskan multimode microplate reader.

For MTT assay, cells were incubated with MTT substrate for 2 h. After extensive washing the resultant formazan crystals were dissolved in DMSO and measured at 595 nm.

### IL1β ELISA

IL1β levels were assayed in cell media using a commercial ELISA kit following manufacturer’s instructions and maximum sensitivity conditions. Absorbance was read at 450 nm in a Varioskan microplate reader.

### ROS detection

For ROS detection, cells were incubated with 10 μM 2′,7′-dichlorofluorescin (DCF) in phenol red-free DMEM and fluorescence was monitored for 30 min at 37°C using a Varioskan plate reader (Ex/Em: 485/535 nm). ROS production was quantified from the slope of the fluorescence increase over time.

### RNA isolation and RT-qPCR

Total RNA was extracted from cells using an RNA isolation kit according to the manufacturer’s instructions. RNA concentration and purity were determined spectrophotometrically by measuring absorbance at 260 and 280 nm. Equal amounts of RNA were reverse transcribed into complementary DNA (cDNA) using a reverse transcription kit (NZY RT master mix) following the manufacturer’s protocol for random hexamers. Quantitative real-time PCR (RT-qPCR) was performed in duplicate using TaqMan Gene Expression Assays and NZYspeedy qPCR Probe Master Mix on a LightCycler^®^ 480 real-time PCR system according to the manufacturer’s instructions. Thermal cycling conditions consisted of an initial activation step at 95°C for 10 min, followed by 40 cycles of denaturation at 95°C for 15 s and annealing/extension at 60°C for 1 min. Relative gene expression was determined using the comparative Ct (2^-ΔΔCt^) method. Expression levels of target genes were normalized to the endogenous control gene 18S. TaqMan assay identifiers used in this study are listed in Reagents Table.

### Phagocytosis assay

Phagocytic activity was determined via internalization of fluorescent Carboxylate-Modified Microspheres (Beads) by confocal microscopy. Cells seeded in 12 mm-coverslips were incubated with BLT-1 for 30 min prior to a 3-hour incubation with the beads in the presence or absence of LPS. Non-internalized beads were washed two times with PBS. Cells were fixed in PFA 4% in PBS for 15 min in the dark. Cell body was visualized using Iba-1 immunostaining while nuclei were stained with DAPI 300 nM and coverslips were mounted with Prolong Diamond mounting media. Images were taken in a LEICA TCS SP8 STED 3X confocal microscope. The number of internalized beads per cell was used as a proxy of phagocytic activity.

### NBD-cholesterol uptake

Cholesterol uptake was determined by internalization of a fluorescent cholesterol analog by fluorimetric measurements or confocal microscopy. Cells were seeded in either 96-well plate or 12 mm-coverslips and incubated with the indicated inhibitors 30 min prior to a 3-hour co-incubation with NBD-cholesterol and LPS. After extensive washing, 96-well plate fluorescence emission (ex/em 485/511) of internalized NBD-cholesterol was measured in a Varioskan microplate reader. For confocal microscopy, cells were washed and fixed with PFA 4% in PBS. Nuclei were stained with DAPI 300 nM for 10 min and coverslips were mounted with Prolong Diamond. Images were taken in a LEICA TCS SP8 STED 3X confocal microscope. NBD-cholesterol fluorescence volume was normalized by the number of cells per image.

### OlyA purification

OlyA-His6 was overexpressed in *E. coli* BL21 and purified as previously described^26^, with minor modifications. The lysis buffer composition was 50 mM Na₃PO₄ pH 7.5, 300 mM NaCl, 0.5% Glycerol, 10 mM Imidazol, 1 mM DTT, 0.05 mg/mL DNase and protease inhibitors. The homogenization was done using a french press and the post-lysis centrifugation was carried out at 25,000×g for 5min. OlyA-His6 was isolated from the lysate using a cobalt column. The eluted OlyA-His6 was then subjected to dialysis at 4°C overnight. Protein labeling was verified by SDS-PAGE followed by Western blotting (antibody anti-His-tag). All purified protein was stored at -20°C in 20% glycerol.

### D4H-mCherry purification

GST-tagged D4H-mCherry fusion protein was overexpressed in *E. coli* C41 and purified as described^52^, with minor modifications, using glutathione Sepharose 4B beads according to the manufacturer’s instructions. Protein purity across different batches was assessed by Coomassie Blue staining. Purified protein was stored at -20°C in the presence of 20% (w/v) sucrose.

### Lipid stainings

Cells were seeded in 12 mm-coverslips and subjected to stated treatments. For sphingomyelin-free cholesterol and sphingomyelin-cholesterol complex detection, cells were respectively incubated with OlyA (5 µg/mL) or DH4-mCherry (50 μg/mL) in PBS supplemented with 1%BSA and TruStain FCX Plus (1:200) for 15 min at r.t. in the dark. After incubation, cells were fixed in 4% PFA in PBS and OlyA-stained cells were further incubated with anti-HisTagAPC (1:100) for 30 min at 4°C in the dark. Free cholesterol and lipid droplet detection was performed post-fixation by incubation with 0.5 mg/mL Filipin Complex III for 2 h or 10 µM BODIPY staining for 15 min at r.t., respectively. For detection of gangliosides-enriched regions in the plasma membrane, cells were incubated with 1 μg/mL cholera toxin B for 15 min at 4°C and then fixed in 4% PFA in PBS.

Nuclei were stained with DAPI 300 nM (except Filipin-stained cells) and coverslips were mounted with Prolong Diamond.

### Image acquisition, analysis and data quantification

All confocal images from a given experiment were acquired using the same laser intensity and photomultiplier settings to avoid variability or bias. Only nuclear DAPI staining was used to select the field to photograph. Images were acquired using a Leica TCS SP8 STED 3X confocal system. Images were processed with Fiji/ImageJ and Imaris 11.0 software.

Unless stated otherwise, maximum projections of all z-stacks are displayed in representative images (z-step: 0.5 µm for cells). For 3D image analysis and segmentation, Imaris 11.0 was utilized. Quantification of OlyA and Cholera toxin B in cells was conducted in 3D using the Imaris Surfaces module through an automated machine learning-based workflow. Total signal volume was extracted from the generated statistics. In regions where cells appeared adjacent or touching, manual mode intervention was applied to ensure accurate individual cell recognition. Representative images display the 3D rendering of the resulting segmentation.

Filipin, BODIPY and NBD-cholesterol quantification was performed using a semi-automatic custom ImageJ/FIJI macro to measure signal volume and number of spots (≥ 3 voxels in size) above a specific intensity threshold. Analysis was restricted to manually defined regions of interest (ROIs), representing either individual cells or entire fields of view normalized by total cell count.

### TLR4 determination in detergent-resistant membranes

An adapted protocol was followed to isolate detergent-resistant membranes^53^. Briefly, cells were washed twice with ice-cold PBS and collected by centrifugation at 100×g for 5 min at 4°C. To lysate cells, the pellet was resuspended and incubated with 5% Triton-X 100 in the extraction buffer (10 mM HEPES, 150 mM NaCl, 5 mM EDTA, pH 7.5) on ice for 1 h. Samples were centrifuged at 12,000×g for 30 min at 4°C and supernatants were collected as the soluble fraction. Pellets were further washed in the extraction buffer and solubilized by sonication in RIPA buffer supplemented with 1 mM EDTA and 1 mM EGTA. After a centrifugation at 12,000×g for 30 min at 4°C, the resulting supernatants were collected as the insoluble fraction. All buffers were supplemented with protease inhibitors.

To detect TLR4 levels in detergent-resistant membrane fractions, equal amounts of protein were loaded and separated in SDS-PAGE and transferred into 0.2 μm PVDF membrane. Non-specific binding sites of the membrane were blocked with 5% skimmed milk in PBS with 0.05% Tween-20 (PBS-T). Primary antibodies for TLR4 and Flotillin-1 were used and incubated (1:1,000 in 3%BSA, 0.02% sodium azide in PBS) overnight at 4°C. Membranes were incubated with the respective anti-HRP secondary antibodies (1:2,000 in 1% BSA in PBS-T) for 1 h at r.t. and detection was performed using the ECL system. Protein bands were quantified by densitometric analysis using ImageJ software.

### Membrane fluidity

Membrane fluidity was assessed using the fluorescent probe pyrene decanoic acid (PDA) which forms excimer upon spatial interaction, shifting the emission maximum from 400 nm to 470 nm. Thus, excimer:monomer ratio (470/400 nm) can be used as a proxy of membrane fluidity. Cells were incubated with 20 μM PDA and 0.08% pluronic F127 in HEPES/Tris buffer (100 mM sodium chloride in 5 mM HEPES/Tris Buffer, pH 7.4 with 2 mM MgCl_2_, 5 mM potassium chloride, 2 mM calcium chloride, 5 mM glucose and 50 mM mannitol) for 20 min at 25° in the dark. Unincorporated PDA was removed by two washes with the buffer solution. Fluorescence was read at 470 and 400 nm with a 360 nm excitation in the pre-warmed Varioskan microplate reader at 37°C. Measures were repeated 5 times in 30-second intervals. Membrane fluidity was expressed as the excimer:monomer (470/400 nm) fluorescence ratio.

### Membrane permeability

The permeability of a membrane to the non-fluorescent fluorescein diacetate (FDA) is driven by passive diffusion. Upon incorporation, FDA is rapidly converted to the fluorescent fluorescein. Therefore, kinetic measurement of fluorescence increases over time immediately after addition of FDA can be used as a proxy of membrane permeability^54^. Briefly, cells were washed with PBS with calcium and magnesium chloride and incubated with 2 μg/mL FDA. Fluorescein fluorescence was measured every 30 seconds for a period of 3 min. The slope of fluorescence increase over time was used as a proxy of membrane permeability. Linearity was kept for the measurement period.

### Determination of cholesterol and cholesterol ester levels

Cholesterol and cholesterol ester levels were assayed using a commercial Amplex® Red kit. Briefly, lipids were extracted from cell pellets or membrane preparations by sequential addition of water, chloroform and methanol (1:2:1), followed by centrifugation at 8,000×g for 5 min. The lower organic phase was collected and dried in a SpeedVac. Fluorescence (Ex/Em 540/590 nm) was measured according to the manufacturer’s instructions using a Varioskan microplate reader at 37°C, with 12 measurements taken at 5-min intervals. Stabilized fluorescence values were used for quantification and normalized to protein content.

### Cholesterol efflux assay

To assess cholesterol efflux, BV2 cells were loaded with 5 μCi/mL ^3^H-cholesterol pre-equilibrated in 2% fatty-acid free BSA (BSA-FFA) for 2h. After extensive washing with PBS, cells were incubated for an additional 4h in 2%BSA-FFA in the presence of 2 μM BLT-1 or 1 μg/mL LPS. Radioactivity released into the medium, reflecting cholesterol efflux, was quantified using a scintillation counter.

### Flow cytometry of BV2 and primary microglia

BV2 or primary microglia were washed twice with PBS and collected by trypsinization (BODIPY, OlyA and D4H-mCherry staining) or scraping in PBS containing 1% BSA (Cholera toxin B and TLR4 staining). Cells were pelleted and stained in Flow Cytometry Buffer (FCB; 0.2% BSA and 2 mM EDTA in PBS) under the conditions described below. Following staining, cells were washed and resuspended in FCB. For viability assessment, 1 μg/mL DAPI was added immediately before acquisition. Samples were analyzed using a CytoFLEX S cytometer and CytExpert software, with viable singlets identified as DAPI^−^ cells. Unstained and assay-specific negative controls were included to establish acquisition parameters and gating.

#### Neutral lipid and lipid raft staining

Neutral lipids were detected by incubating cells with 10 μM BODIPY for 15 min at room temperature. GM1, used as a proxy for lipid raft content, was detected by incubating cells with fluorescently labeled Cholera toxin B (1 μg/mL) in 1% BSA for 30 min at 4°C.

#### SM/cholesterol determination by OlyA

Cells were incubated with OlyA (25 μg/mL) and TruStain FCX Plus (1:200) in FCB for 30 min at 4°C. After washing, cells were incubated with anti-HisTag-APC (1:100) in FCB for 30 min at 4°C in the dark, followed by two washes. OlyA binding was detected in the APC channel. Negative controls lacking OlyA or anti-HisTag-APC were included for gating.

#### Accessible cholesterol detection by D4H-mCherry

Cells were incubated with D4H-mCherry (25 μg/mL) and TruStain FCX Plus (1:200) in FCB for 15 min at 4°C. Cells were then fixed with 4% PFA in PBS for 10 min at room temperature in the dark, washed, and resuspended in PBS. D4H-mCherry fluorescence was measured using the mCherry-PI-TR channel.

#### TLR4 endocytosis

Cells were incubated with TruStain FCX Plus (1:200) for 10 min on ice, followed by TLR4-PE antibody (1:800) for 15 min on ice in the dark. Cells were washed twice and TLR4 fluorescence was measured in the PE-DsRed channel. A PE Rat IgG2a control was included to assess nonspecific binding. A decrease in PE mean fluorescence intensity (MFI) among viable cells over time was used as a proxy for TLR4 endocytosis.

### Flow cytometry for adult murine microglia

#### Single cell suspension preparation

Single cell suspension from murine brain was prepared as previously described^55^. Briefly, mice were sacrificed via cervical dislocation and brains were immediately extracted and rinsed in ice-cold MEM. One hemisphere per condition was used. All steps were carried out on ice. After removing the cerebellum, the remaining brain tissue was minced and mechanically dissociated using a Dounce homogenizer (around 15-20 strokes with pestle A followed by 10 strokes with pestle B) until a homogeneous suspension was achieved. The homogenate was filtered through a pre-wetted 70 μm strainer and centrifuged at 300×g for 5 min. To remove myelin debris, a two phase solution was created by resuspending the cell pellet in a 25% isotonic percoll solution (3:1 PBS:percoll) and by carefully adding an equivalent volume of PBS on top. After a 10 min centrifugation at 300×g and 4°C, the supernatant was carefully removed and the resulting cell pellet was washed with PBS and transferred into pre-blocked (5% BSA) cytometer tubes for downstream stainings.

#### SM/cholesterol determination by OlyA

SM/cholesterol complexes in the plasma membrane of microglia (Tmem119-GFP^+^) were analyzed by OlyA-His6 detection by flow cytometry. Cells were incubated with OlyA-His6 (50 μg/mL) and TruStain FCX Plus (1:200) in FCB for 30 min at 4°C. After incubation, cells were washed with FCB and incubated with anti-HisTagAPC (1:100) for 30 min at 4°C in the dark. Cells were pellet and washed twice with FCB buffer and resuspended in 1 μg/mL DAPI in FCB. Samples were flown through the CytoFlexS and OlyA-APC was analyzed in the viable (DAPI^-^) microglia (GFP^+^) population. Negative controls for OlyA and HisTag APC were included to set acquisition parameters and gates.

#### Accessible cholesterol determination by D4H-mCherry

The accessible pool of cholesterol (not sequestered by sphingomyelin) in the plasma membrane was detected by D4H-mCherry sensor. Briefly, cells were incubated with DH4-cherry (500 μg/mL) and TruStain FCX Plus (1:200) in FCB. After a 15 min incubation at 4°C, cells were washed with PBS and resuspended in FCB containing 1 μg/mL of DAPI. Samples were flown and analysed in the CytoFlexS using mCherry-PI-TR (D4H-mCherry detection), FITC-GFP (Tmem119-GFP microglia detection) and PB450-DAPI (cell viability) detection channels.

### Targeted Lipidomics and Data analysis

Lipid extraction was performed following the procedure previously described^56^. In brief, 10 µL of sample were mixed with 100 µL of a 1:1 (v/v) butanol:methanol solution supplemented with 10 mM ammonium formate and the corresponding internal standards (ISTD, Reagents Table). Samples were vigorously vortexed and subsequently sonicated for 1 h at room temperature. After sonication, extracts were centrifuged at 14,000 × g for 10 min at 20°C. The resulting supernatant was collected and transferred into glass-insert vials for subsequent lipidomic analysis.

Lipid profiling was carried out using an Ultivo Triple Quadrupole mass spectrometer coupled to an Agilent 1290 UHPLC system. Chromatographic separation was achieved on a ZORBAX Eclipse Plus C18 column (2.1 × 100 mm, 1.8 µm) maintained at 60°C. Data acquisition was performed in both positive and negative electrospray ionization modes using dynamic scheduled multiple reaction monitoring.

The mobile phase consisted of solvent A, containing H₂O/acetonitrile/isopropanol (50:30:20, v/v/v) with 10 mM ammonium formate, and solvent B, composed of H₂O/acetonitrile/isopropanol (1:9:90, v/v/v) supplemented with 10 mM ammonium formate. Separation was performed at an initial flow rate of 0.4 mL/min with 10% solvent B. The gradient increased to 45% B within 2.7 min, followed by sequential increases to 53% B (0.1 min), 65% B (6.2 min), 89% B (0.1 min), 92% B (1.9 min), and finally 100% B (0.1 min). This composition was maintained for 2.3 min, resulting in a chromatographic run time of 13.4 min. Re-equilibration was achieved by returning to 10% B over 0.1 min and holding this composition for 0.9 min. During this step, the flow rate was increased to 0.6 mL/min for 1 min and subsequently reduced to 0.4 mL/min over 0.1 min. The initial mobile-phase conditions were maintained for an additional 0.9 min, giving a total cycle time of 16.5 min.

Lipid species were identified based on their characteristic precursor-to-product ion transitions and retention times, as previously reported^57^. Chromatographic peak integration was performed using Agilent MassHunter Quantitative Analysis software. Relative quantification was obtained by normalizing the peak area of each analyte to that of its corresponding internal standard.

All statistical analyses were performed in R 4.4.2, using the packages and tools specified in each methodological section.

#### Annotation and preprocessing

Lipid annotations provided by the analytical platform were standardized using RefMet^58^. Entries that could not be unambiguously interpreted were identified and manually curated before repeating the conversion. These included diacylglycerols carrying additional acyl-chain annotations, TAG species reported through precursor-ion ([SIM]) or neutral-loss ([NL-x:y]) transitions, isomer-specific neutral-loss annotations, ether-linked TAG (O-TG), and cholesterol entries originally reported as CHO. Final assignments were cross-checked against structural information provided by the analytical platform and reference resources, including LIPID MAPS^59^ and Metabolomics Workbench^60^. The final analytical panel comprised 335 lipid species, including 269 structural membrane lipids (197 glycerophospholipids and 72 sphingolipids) and 66 storage lipids. Sixteen missing observations distributed across ten lipid species were retained without imputation. Lipid abundances were normalized using probabilistic quotient normalization (PQN)^61^ based on a reference set of structural membrane lipids and free cholesterol. Normalized values were subsequently log2-transformed prior to statistical analysis.

#### Differential abundance analysis

Differential abundance was assessed using limma v3.62.2^62^. A cell-means linear model (∼ 0 + group) was fitted to the normalized log2-transformed abundance matrix, and the following contrasts were extracted from a single model: BLT-1 vs Control, LPS vs Control, LPS + BLT-1 vs Control, LPS + BLT-1 vs LPS. P values were adjusted for multiple testing using the Benjamini-Hochberg false discovery rate (FDR) procedure. Lipid species were considered differentially abundant at FDR < 0.05.

#### Subclass-level analysis

Subclass-level changes were assessed using fry, implemented in limma v3.62.2^63^. Effect sizes were estimated from subclass-level summaries obtained by centering individual lipid species and calculating the median value per subclass and sample. Only subclasses containing at least three lipid species were included.

The relative contribution of LPS, BLT-1, their interaction, and residual variation was additionally estimated for each lipid species using a two-way factorial model (∼LPS*BLT). Variance components were expressed as percentages and averaged within lipid subclasses containing at least three species. This analysis was used descriptively and was not interpreted as a formal significance test.

Relationships between metabolically connected sphingolipid classes were evaluated using three ratios: Cer/SM, Hex2Cer/HexCer, and HexCer/Cer. Ratios were calculated for each sample as the difference between the corresponding subclass medians in log2 space. Group differences were assessed across the four principal contrasts using two-sample t tests with Benjamini-Hochberg correction within each contrast.

#### Lipid reaction network analysis

Sphingolipid reaction networks were generated using LINEX2^64^, which constructs species-level lipid reaction networks based on curated biochemical reaction rules and identifies subnetworks in which substrate-product relationships change between experimental groups. The 72 sphingolipid species included in the final analytical panel were submitted to LINEX2, and network enrichment was evaluated independently for each contrast. The analysis was performed using linear-scale abundances and ratio-based changes, with subnetwork sizes restricted to three to five reactions. When both optimal and suboptimal enriched subnetworks were returned, both were retained to avoid interpreting non-top-ranked reactions as unaffected.

LINEX2 results were visualized using three complementary representations. The first organizes sphingolipid species according to their total carbon number, allowing identification of carbon-length ranges in which differential changes are concentrated. A second representation incorporates both total carbon number and number of double bonds, distinguishing species with identical carbon number but different degrees of unsaturation. The conventional LINEX network representation was additionally used to place the observed changes within metabolic connections between SM, Cer, HexCer, and Hex2Cer and their associated reactions. These network representations were used to contextualize differential abundance patterns and do not constitute direct measurements of metabolic flux or enzymatic activity.

#### Lipid ontology enrichment

Functional lipid enrichment was performed using the LION ontology^65^ obtained through the topOnto.LION.db. Panel identifiers were converted to LIPID MAPS-compatible nomenclature and mapped to ontology terms using molecular-species annotation whenever available and sum composition otherwise.

Lipid species lacking an appropriate automated ontology assignment were manually annotated using existing LION terms according to their known chemical structure. Manual curation included acylcarnitines, ether-linked lysophospholipids, plasmalogens, sphingosine, ubiquinone, 1-deoxyceramides, and dihydroceramides. In particular, 1-deoxyceramides and dihydroceramides were distinguished from canonical N-acylsphingosines and assigned to their appropriate sphingoid-base and ceramide subclasses. Acylcarnitines were annotated according to their mitochondrial association and acyl-chain characteristics rather than treated as free fatty acids.

Enrichment analysis was performed using fgsea v1.32.4 on lipid species pre-ranked according to their log2 fold change for each contrast. Ontology terms containing fewer than four measured species were excluded. P values were adjusted using the BH procedure independently within each contrast, and terms with FDR < 0.05 were considered significantly enriched. Normalized enrichment scores (NES) were used to determine enrichment direction, allowing positively and negatively enriched categories to be evaluated within a single ranked analysis. Because the LION ontology is hierarchical, related significant terms can be highly redundant. Redundant terms were therefore collapsed independently within each contrast. When terms belonged to the same ontology branch, showed the same enrichment direction, and shared at least 80% of their species sets, the most statistically significant term was retained.

### Statistical analysis

Data are presented as mean ± SEM, with each point representing an individual replicate. All averages were obtained from three or more independent experiments performed at different times using independent sets of samples. Data were assumed to follow a normal distribution. Statistical analyses were performed using GraphPad Prism v9.01. The Brown-Forsythe test was used to assess homogeneity of variance between groups. Statistical significance was assessed using two-tailed unpaired *t*-tests, one-way ANOVA, or (repeated-measures) two-way ANOVA, as appropriate, followed by Tukey’s or Šídák’s multiple-comparison post hoc test. Welch’s correction was applied to *t*-tests when a significant difference in variance between groups was detected. The Kolmogorov-Smirnov test was used to compare cumulative distributions between conditions. A *p* value < 0.05 was considered statistically significant. Statistical significance is indicated as follows: *p* < 0.05, \**p* < 0.01, \*\**p* < 0.001, and \*\*\**p* < 0.0001. Investigators were not blinded during the quantification of imaging experiments.

## Data Availability

The lipidomics data is available in Zenodo (#). All reagents used in this work are commercially available, with manufacturer catalog numbers included in the reagents table. Source data is also provided.

## Acknowledgements.

This work was supported by the Spanish Ministry of Science, Innovation and Universities and Agencia Estatal de Investigación (PID2024-161283OA-I00 to JM; FPU fellowship FPU23/02047 to CA-B, FPI fellowship PREP2024-002719 to DG-D; FPU fellowship 22/00245 to MU; RYC2024-048419-I to IA-F; PID2022-139936OA-I00 to EN); Comunidad de Madrid (2024-T1SAL-GL-31373 Talento César Nombela to JM); the European Union’s Horizon 2020 research and innovation programme under the Marie Skłodowska-Curie grant agreement (MSCA-PF 101106857 to JM). JMBP was supported by the Minister of Science and Technology of Colombia. CG-R and FG-G were supported and partially funded by CIAICO/2023/149 and CIGE/2024/242 from the Consellería de Educación, Cultura y Universidades de la Generalitat Valenciana; by PID2021-124430OA-I00, and PID2025-174324OB-I00 from MICIU/AEI/10.13039/501100011033, and ERDF/EU.

We thank Ignacio Ramírez-Pardo for developing the semi-automatic custom ImageJ/FIJI macro and Carla Sevilla-Navarro for helpful advice on the production of OlyA protein. We also thank the Animal Facility, Confocal Microscopy Facility, and Cytometry Core Facility at CIB-Margarita Salas for their technical support.

## Author Contributions Statement

Conceived the project: JM. Designed experiments: JM. Generated experimental data: CA-B, BV-Z, SC-L, JMB-P, DG-D, MU and EN. Collected/analyzed lipidomics data: FB-A, MP-O, CG-R and FG-G. Wrote the manuscript: CA-B and JM. Critically edited the manuscript: all authors. Approved final version of the manuscript: all authors.

## Figure Legends

**Figure S1.**
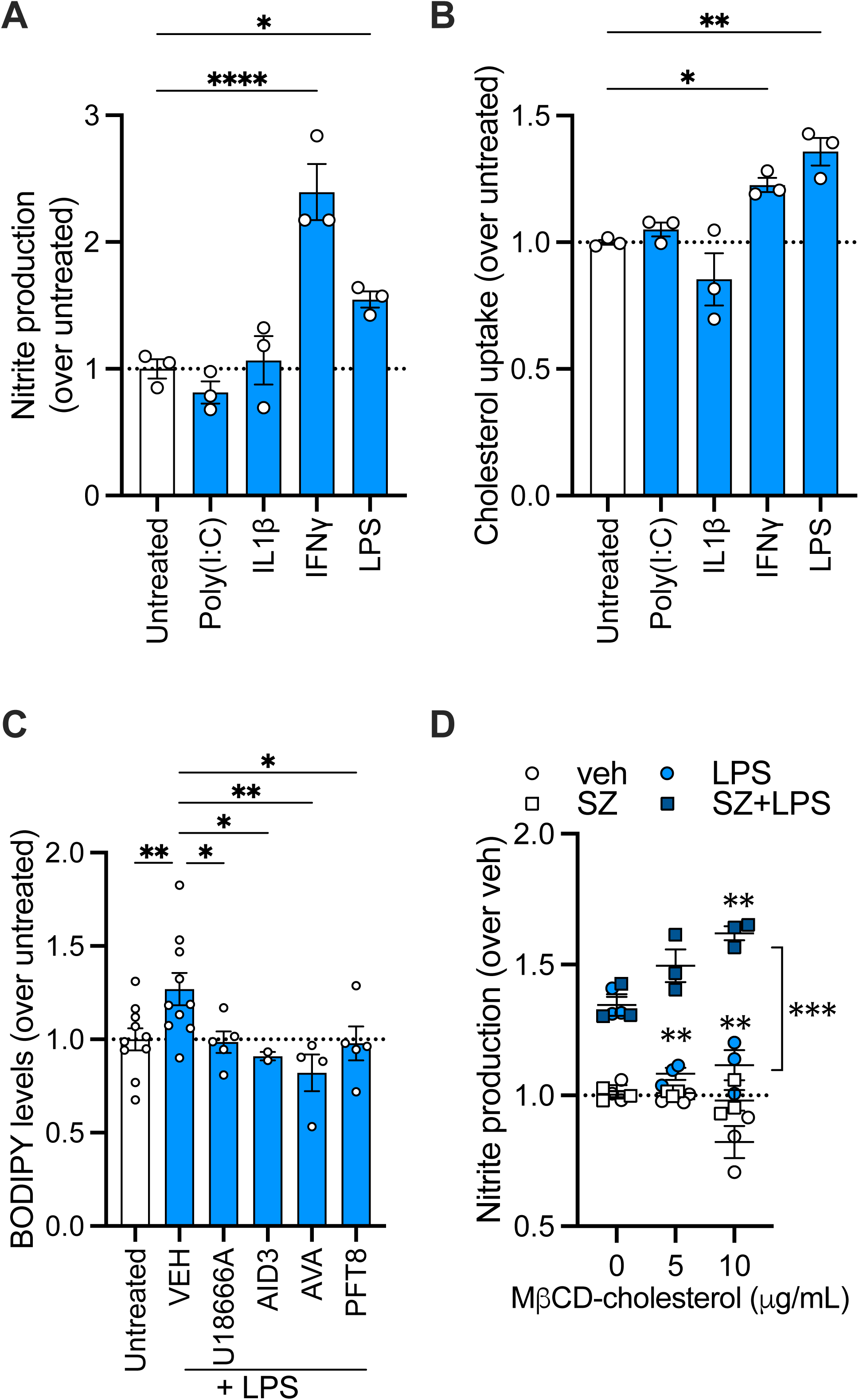
Cholesterol trafficking and esterification modulate inflammatory responses in microglia. **A-B.** BV2 cells were stimulated with Poly(I:C), IL1β, IFNγ, or LPS for 18h, and nitrite production (**A**) and NBD-cholesterol uptake (**B**) were determined. **C.** Lipid droplet levels were quantified by BODIPY staining and flow cytometry in BV2 cells stimulated with LPS for 18h in the presence of U18666A, AID3, the SOAT1 inhibitor avasimibe (AVA), or a combination of DGAT1 and DGAT2 inhibitors (PFT8). **D.** BV2 cells were incubated with increasing concentrations of MβCD-cholesterol for 24h in the presence or absence of the SOAT1 inhibitor Sandoz 58-035 (SZ), followed by LPS stimulation and measurement of nitrite production.

**Figure S2.**
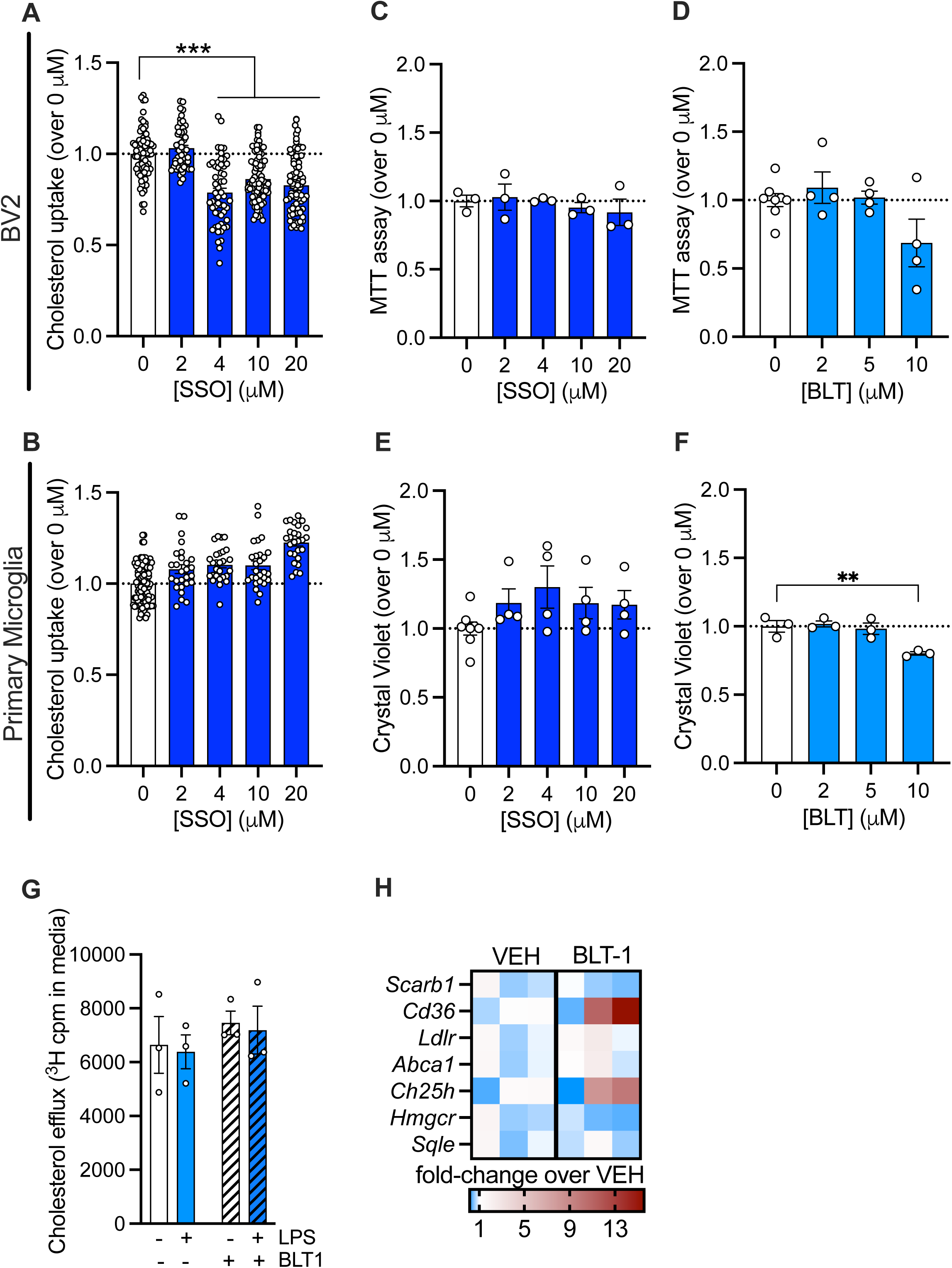
SR-B1 inhibition with BLT-1 and CD36 inhibition with SSO exert distinct effects in BV2 cells and primary microglia. **A-B.** BV2 cells (**A**) or primary microglia (**B**) were treated with increasing concentrations of SSO, and NBD-cholesterol uptake was monitored after 3h. **C-D.** Cell viability was assessed by MTT assay in BV2 cells treated for 18h with the indicated concentrations of SSO (**C**) or BLT-1 (**D**). **E-F.** Primary microglial viability after 18h of treatment with increasing concentrations of SSO (**E**) and BLT-1 (**F**) was evaluated by crystal violet staining. **G.** BV2 cells were loaded with ^3^H-cholesterol (5 µCi/mL) for 2h. After extensive washing, cells were incubated in 2% BSA-FFA in the presence or absence of BLT-1 and LPS, and 4h later, efflux of ^3^H-cholesterol into the medium was analyzed. **H.** Expression of the indicated genes was analyzed by RT-qPCR in BV2 cells treated with BLT-1 for 18h. Data are presented as fold change relative to VEH.

**Figure S3.**
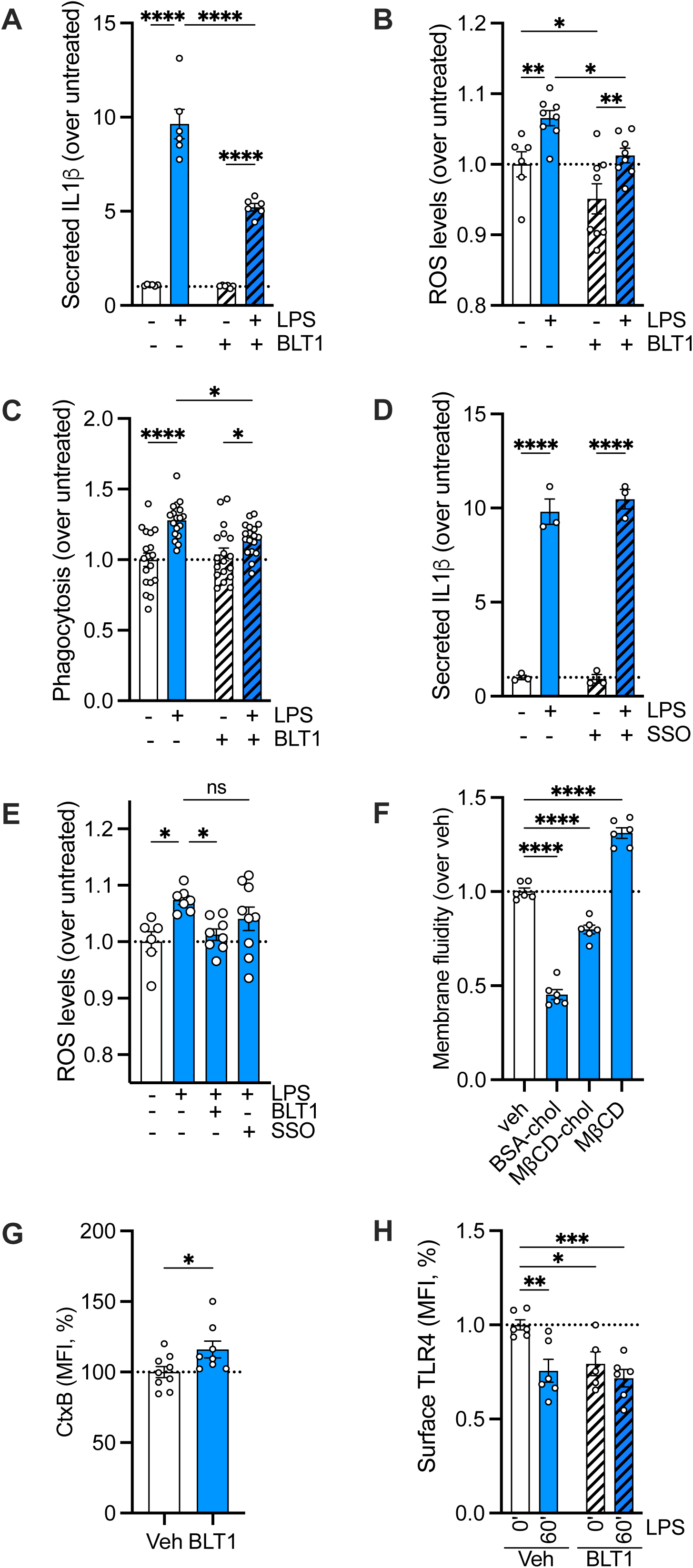
BLT-1 inhibits TLR4-mediated inflammatory responses in primary microglia. **A-C** Primary microglia were incubated with BLT-1 for 18h and stimulated with LPS for 24h. Secreted IL1β (**A**), ROS production (**B**) and phagocytosis (**C**) were subsequently evaluated. **D.** IL1β secretion was measured in primary microglia stimulated with LPS after pre-treatment with 5 µM SSO for 18h. **E.** ROS production was analyzed in primary microglia stimulated with LPS following treatment with BLT-1 or SSO for 18h. **F.** Membrane fluidity was assessed in BV2 cells treated with 25 µg/mL BSA-cholesterol for 18h, 5 µg/mL MβCD-cholesterol for 3h, or 50 µM MβCD for 1 h. **G.** Cholera toxin B staining was analyzed in primary microglia after 18h incubation with BLT-1. **H.** Surface TLR4 levels were determined in primary microglia pre-treated with BLT-1 for 18h and stimulated with LPS for the indicated times.

**Figure S4.**
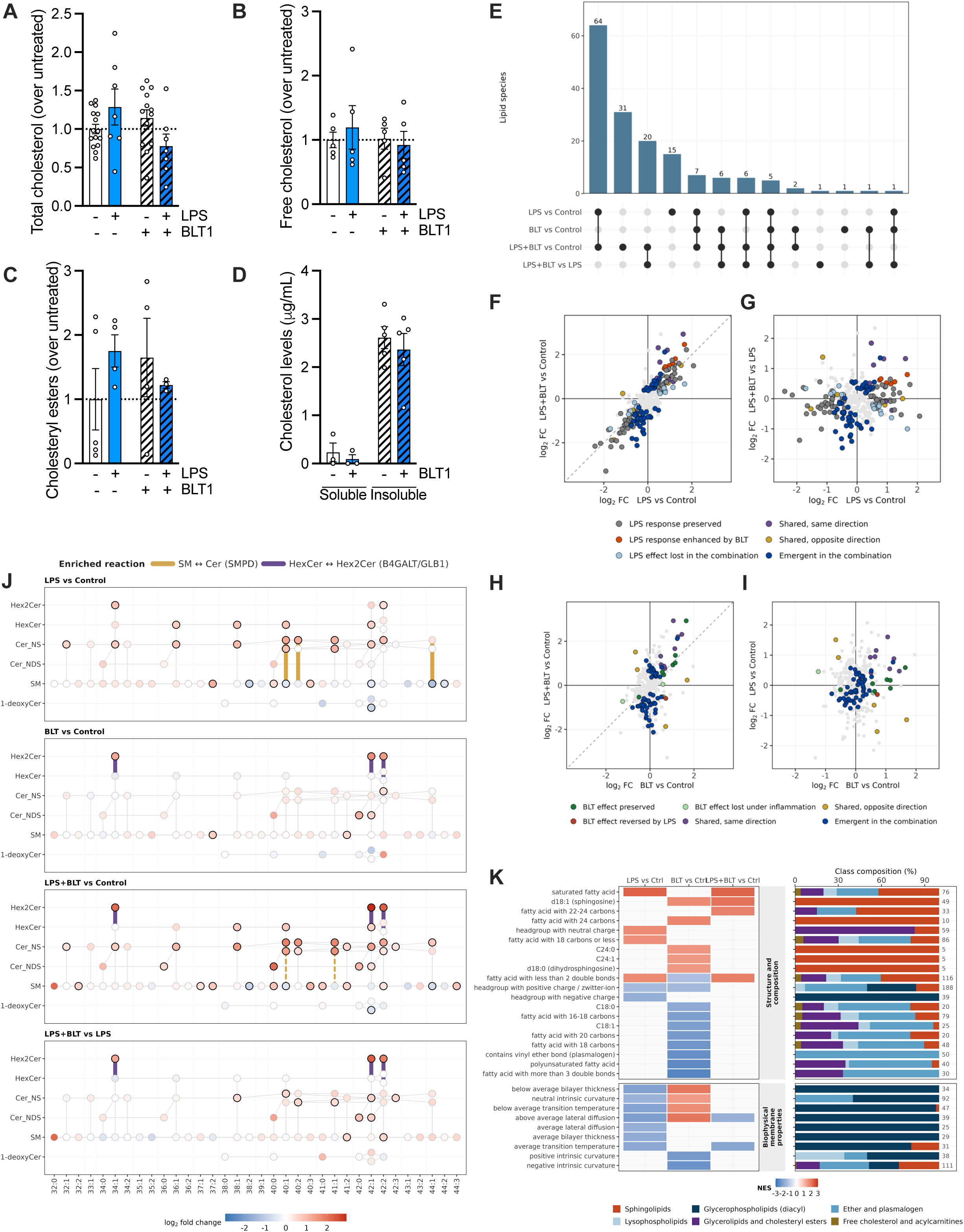
Lipidomic composition is modulated by LPS and BLT-1 treatment. **A-C.** Lipid extracts from BV2 cells pre-treated with BLT-1 for 18h and stimulated with LPS for 24h were analyzed to determine total cholesterol (**A**), free cholesterol (**B**) and cholesteryl ester levels (**C**). **D.** Cholesterol content in Triton X-100-soluble and insoluble fractions from BV2 cells treated with BLT-1 for 18h. **E.** Overlap of differentially abundant lipid species across the four analyzed contrasts. The UpSet plot shows the number of lipid species shared among or specific to LPS vs Control, BLT vs Control, LPS+BLT vs Control, and LPS+BLT vs LPS. Bars indicate the size of each intersection, while the connected dots in the lower matrix identify the contrasts included in each intersection. **F-G.** Modulation of the LPS-induced lipidomic response by BLT-1 pretreatment. (**F**) Comparison of log2 fold changes between LPS vs Control and LPS+BLT vs Control, showing the extent to which the lipidomic response to LPS is preserved or modified following BLT-1 pretreatment. The dashed diagonal indicates equal effect sizes in both contrasts. (**G**) Relationship between the effect induced by LPS (LPS vs Control) and the additional effect associated with BLT-1 under inflammatory conditions (LPS+BLT vs LPS). Lipid species are classified according to their response pattern across contrasts. **H-I.** Context dependence of the lipidomic response to BLT-1. (**H**) Comparison of log2 fold changes between BLT vs Control and LPS+BLT vs Control, showing how lipid species responding to BLT-1 behave in the combined condition. The dashed diagonal indicates equal effect sizes in both contrasts. (**I**) Relationship between the basal effect of BLT-1 (BLT vs Control) and the lipidomic response to LPS (LPS vs Control). Lipid species are classified according to whether the BLT-1-associated response is preserved, lost or modified in the inflammatory context. **J.** Structural distribution of sphingolipid changes across treatments. Sphingolipid species are arranged by subclass and total molecular composition, expressed as total carbon atoms:number of double bonds. Node color represents the log2 fold change for each experimental contrast. Highlighted connections indicate enriched LINEX reactions between SM and Cer (SMPD-associated) and between HexCer and Hex2Cer (B4GALT/ GLB1-associated). **K.** Tile plot showing selected LION enrichment terms related to lipid structure/composition and predicted membrane biophysical properties across the LPS vs Control, BLT vs Control and LPS+BLT vs Control contrasts. Tile color represents the normalized enrichment score (NES), with red indicating positive enrichment and blue indicating negative enrichment. The stacked bars on the right show the lipid class composition of each LION term, expressed as percentage of contributing lipid species. Numbers indicate the total number of lipid species annotated within each term. This representation links enriched structural and membrane-related features with the lipid classes driving each signal.

**Figure S5.**
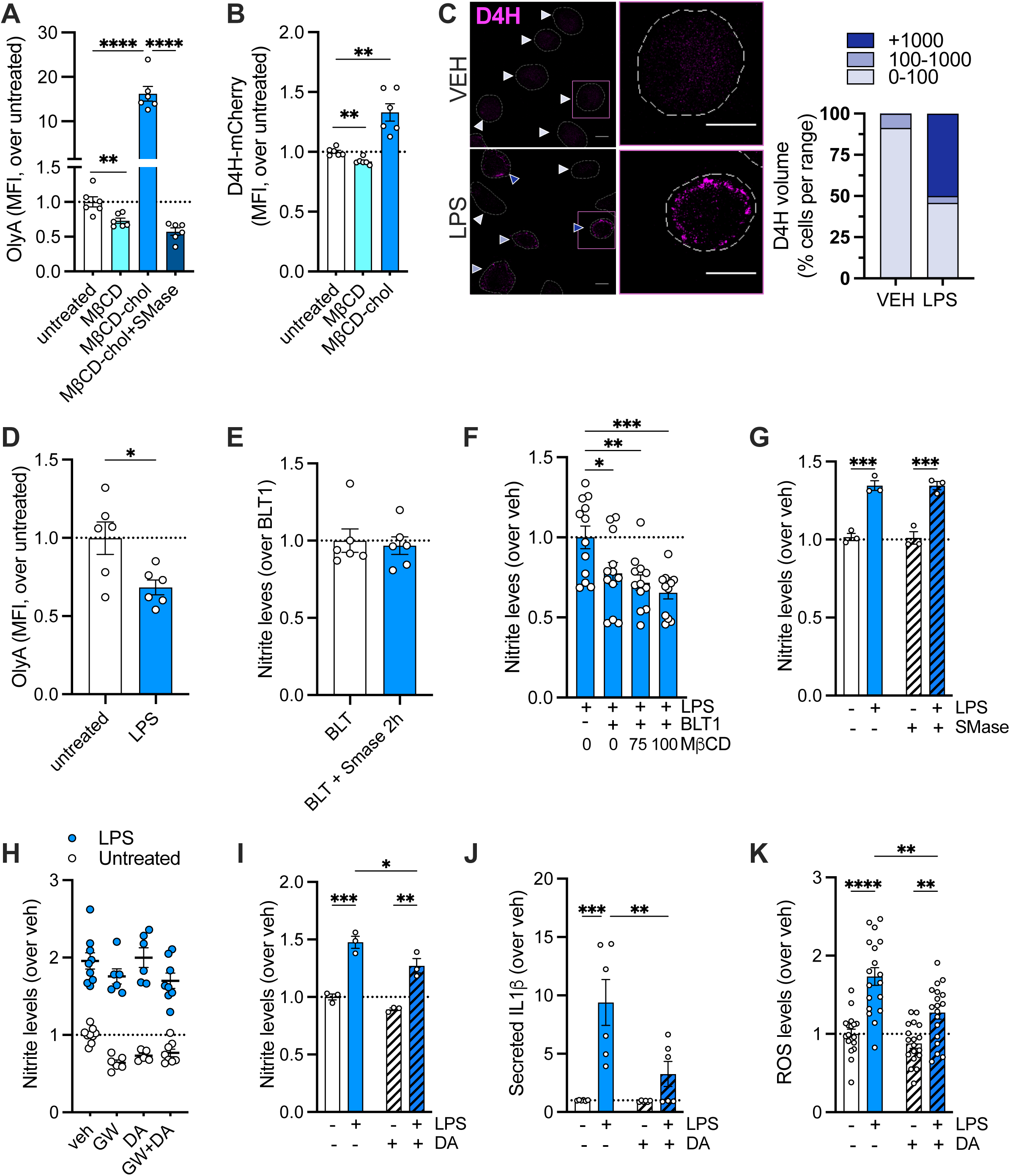
The balance between accessible and inaccessible cholesterol pools regulate LPS-induced inflammatory signaling. **A.** OlyA staining was analyzed in BV2 cells treated with 200 µM MβCD, 60 µg/mL MβCD-cholesterol and 60 µg/mL MβCD-cholesterol plus 0,5 µl/mL of SMase for 1h. **B.** D4H-mCherry fluorescence was measured by flow cytometry in BV2 cells treated with 200 µM MβCD or 60 µg/mL MβCD-cholesterol for 1h. **C.** Confocal analysis of D4H-mCherry of primary microglia treated with LPS for 18h. D4H signal was acquired by confocal microscopy and quantified to obtain D4H^+^ volume per cell. Individual cells were then binned according to D4H volume (0-100, 100-1000, and >1000 arbitrary volume units). The proportion of cells within each volume category was calculated and expressed as a percentage of total cells per condition. Distribution shifts between conditions were further assessed using cumulative distribution analysis and the Kolmogorov-Smirnov test. Scale bar = 10 µm. **D.** OlyA staining was used to assess the inaccessible cholesterol pool in primary microglia stimulated with LPS for 18h. **E.** BV2 cells were pre-treated with BLT-1 for 18h, incubated with 0,5U/mL SMase for 2h, and subsequently stimulated with LPS. Nitrite production was then measured. **F.** BV2 cells were pre-treated with BLT-1 for 18h, followed by incubation with 75-100 µM MβCD for 2h before LPS stimulation. Nitrite production was subsequently analyzed. **G.** BV2 cells were treated with 0,5 U/mL SMase for 18h and stimulated with LPS for 24h. Nitrite production was then quantified. **H.** BV2 cells were stimulated with LPS for 24h in the presence of the neutral SMase inhibitor GW4869 (GW) or the acid SMase inhibitor desipramine (DA), and nitrite production was analyzed. **I-K.** Primary microglia were pre-treated with desipramine for 18h followed by LPS stimulation. Subsequently, nitrite production (**I**), IL1β secretion (**J**) and ROS levels (**K**) were measured.

**Table S1** - Lipidomic composition of BV2 cells pre-treated with BLT-1 for 18h and stimulated with LPS for 24h.

**Table S2** - Differential abundance of significant lipid species detected.

**Table S3** - Representative LINEX-enriched sphingolipid reactions across experimental contrasts.

