## Supplemental Table 3 for "A sphingomyelin-cholesterol complex regulates TLR4 activation in microglia"

**Table S3 -** *Representative LINEX-enriched sphingolipid reactions across experimental contrasts. The table shows representative significantly enriched LINEX-inferred reactions involving sphingolipid species across 10 independent runs. For each reaction, the associated enzyme candidates, enrichment score, p value and contributing lipid species are shown. Reactions should be interpreted as pathway-level associations between compositionally compatible lipid species rather than definitive one-to-one molecular conversions, particularly for sphingomyelin species annotated at sum-composition level. ENPP7, alkaline sphingomyelinase/ectonucleotide pyrophosphatase-phosphodiesterase 7; ASM, acid sphingomyelinase; SMPD1-4, sphingomyelin phosphodiesterases 1–4; GLB1, β-galactosidase 1; B4GALT1/3, β-1,4-galactosyltransferases 1/3; REQ_38260, LINEX reaction identifier; SM, sphingomyelin; Cer, ceramide; HexCer, hexosylceramide; Hex2Cer, dihexosylceramide.*

| **CONTRAST** | **REACTION** | **ENZYMES** | **SCORE** | **P VALUE** | **SPECIES** |
| --- | --- | --- | --- | --- | --- |
| LPS vs Control | SM ↔ Cer | ENPP7, ASM-, SMPD1/2/3/4 | 1.99 | 0.0074 | SM 40:1;O2; SM 40:2;O2; SM 44:1;O2; Cer 18:1;O2/22:0; Cer 18:2;O2/22:0; Cer 20:1;O2/24:0 |
| BLT vs Control | Hex2Cer ↔ HexCer | GLB1, B4GALT1, B4GALT3 | 2.11 | 0.0053 | HexCer 18:1;O2/16:0; HexCer 18:1;O2/24:0; HexCer 18:1;O2/24:1; Hex2Cer 18:1;O2/16:0; Hex2Cer 18:1;O2/24:0; Hex2Cer 18:1;O2/24:1 |
| LPS+BLT vs Control | Hex2Cer ↔ HexCer | GLB1, B4GALT1, B4GALT3 | 3.95 | 0.013 | HexCer 18:1;O2/16:0; HexCer 18:1;O2/24:0; HexCer 18:1;O2/24:1; Hex2Cer 18:1;O2/16:0; Hex2Cer 18:1;O2/24:0; Hex2Cer 18:1;O2/24:1 |
| LPS+BLT vs Control | SM ↔ Cer | ENPP7, ASM-, SMPD1/2/3/4 | 3.51 | 0.0252 | SM 40:1;O2; SM 41:1;O2; Cer 18:1;O2/22:0; Cer 18:1;O2/23:0; Cer 17:1;O2/24:0 |
| LPS+BLT vs LPS | Hex2Cer ↔ HexCer | GLB1, B4GALT1/3; REQ_38260 | 3.14 | 0.007 | Cer 18:1;O2/24:0; Cer 18:1;O2/24:1; HexCer 18:1;O2/16:0; HexCer 18:1;O2/24:0; HexCer 18:1;O2/24:1; Hex2Cer 18:1;O2/16:0; Hex2Cer 18:1;O2/24:0; Hex2Cer 18:1;O2/24:1 |
